# HTLV-1-Induced Neuroimmunome Correlates with Disease Progression and Severity

**DOI:** 10.64898/2026.03.12.711373

**Authors:** Fernando Yuri Nery do Vale, Carlota Miranda-Solé, Adriel Leal Nóbile, Júlia Nakanishi Usuda, Dennyson Leandro M Fonseca, Lena F. Schimke, Mauro Cesar Cafundó Morais, Débora Gomes de Albuquerque Freitas, Anny Silva Adri, Roseane Galdioli Nava, Yohan Lucas Gonçalves Corrêa, Hugo Fernando Nery do Vale, Adriana Simizo, Niels Olsen Saraiva Câmara, Gustavo Cabral-Miranda, Helder I. Nakaya, Rodrigo J. S. Dalmolin, Haroldo Dutra Dias, Yuki Saito, Yasunori Kogure, Junji Koya, Keisuke Kataoka, Igor Salerno Filgueiras, Margarita Dominguez-Villar, Otávio Cabral Marques

**Author notes:** Corresponding authors: Fernando Yuri Nery do Vale;, Otavio Cabral-Marques, MSc, PhD. Contributed Equally (Fernando Yuri Nery do Vale, Carlota Miranda-Solé, Margarita Dominguez-Villar).

## Abstract

Human T-cell lymphotropic virus type 1 (HTLV-1) is a retrovirus that infects approximately 5–10 million people worldwide. While most individuals remain asymptomatic, a significant subset develops debilitating neuroinflammatory or malignant disorders, including adult T-cell leukemia/lymphoma (ATL) and HTLV-1-associated myelopathy/tropical spastic paraparesis (HAM/TSP). To unravel the systemic molecular mechanisms underlying HTLV-1 pathogenesis, we employed a multi-dimensional systems biology approach, integrating bulk transcriptomic data from total PBMCs (n = 200) with single-cell RNA sequencing (scRNA-seq) from 233,093 peripheral blood mononuclear cells (PBMCs). Our analysis revealed a consistent and clinically relevant neuroimmune signature within leukocytes, termed the neuroimmunome, comprising a set of differentially expressed genes shared across the nervous and immune systems. Through dimensionality reduction and machine learning techniques, such as PCA, gradient boosting, and MANOVA with bootstrapping, we identified potential biomarkers predictive of HTLV-1-driven leukemogenesis, which were subsequently validated across ATL, HAM/TSP, and asymptomatic cohorts via flow cytometry. Notably, expression levels of proteins such as ATF4 and SKIL were strongly correlated with proviral load, suggesting that sustained neuroimmune dysregulation may contribute to disease progression. These findings highlight a previously underappreciated neuroimmunological layer, redefining HTLV-1-associated disease as a condition deeply rooted in neuroimmune network disruption within leukocytes and offering potential novel targets.

## INTRODUCTION

Human T-cell leukemia virus type 1 (HTLV-1) integrates its genome into peripheral blood mononuclear cells (PBMCs), predominantly CD4+ T cells, resulting in a chronic, persistent infection that often remains asymptomatic for decades^1,2^. However, an estimated 5 million to 10 million people worldwide are infected with the virus^3^. Approximately 5% of infected individuals develop adult T-cell leukemia/lymphoma (ATL)^1^, while around 4% develop HTLV-1-associated myelopathy/tropical spastic paraparesis^4^ (HAM/TSP). A growing body of evidence highlights the importance of neuroimmune crosstalk in various disease conditions, including infectious diseases^5,6^, cancer^7–9^ and autoimmune diseases^10–12^. Considering that patients infected with HTLV-1 present with neuroinflammation, synaptic alterations, and microglial cell activation, this suggests that neuroimmune interactions are involved in the disease’s immunopathogenesis^13–15^.

The neuroimmune interaction is a complex network involving various cells from both the immune and nervous systems^16,17^. It can be modulated by different factors, including cytokines^18,19^, neurotransmitters^20,21^, and synaptic signaling^22,23^. In this context, various immune components, such as leukocytes and lymphoid organs, can be activated and influenced by neuroimmune interactions^24–27^. For example, both immune cells and neurons can produce and respond to neurotransmitters and cytokines through specific receptors^18,28^. This suggests that not only neurons, but also immune cells possess sophisticated molecular machinery, including synaptic molecular networks and other genes that regulate neuronal biological processes.

Recent advances in neuroimmunology suggest that circulating immune cells are carriers; rather, they, but may express a range of synaptic molecules and neurotransmitter receptors, participating in bidirectional signaling with the central nervous system^28^. This insight has led us to define the “neuroimmunome”, a set of neuroimmune-related genes (gene clusters) that are intrinsically present in leukocytes^29,30^. Whether these gene signatures are associated with the diverse clinical outcomes of HTLV-1 infection, and whether they underlie mechanistic links between immune surveillance, neurodegeneration, and leukemogenesis, remains to be elucidated.

To explore this hypothesis, we conducted a systems-level transcriptomic analysis of PBMCs from patients with ATL, asymptomatic HTLV-1 carriers (ACs), HAM patients, and healthy donors (HD). Our goal was to identify the neuroimmunome signatures that distinguish disease phenotypes, offering new insights into the molecular underpinnings of HTLV-1-induced clinical diversity.

## MATERIAL AND METHODS

Detailed materials and methods are available in the Supplementary Data.

## Results

### scRNA-seq profiling reveals cell-type diversity and malignant expansion in HTLV-1 infection

To understand how neuroimmune alterations manifest across the clinical spectrum of HTLV-1 infection, we examined peripheral blood mononuclear cell (PBMC) populations at single-cell resolution (**Supplementary Table 1**). Our cohort included 11 ACs (11 samples), 30 patients with ATL, comprising 19 acute, 12 chronic, and 3 smoldering cases, and 3 HD.

Unsupervised clustering of transcriptomic profiles revealed five distinct immune cell populations, including B cells, myeloid cells, natural killer cells, T_h_ CD4+, and T_c_ CD8+ cells **(Fig.1a** and **Suppl. Fig. 2a**), reflecting the cellular complexity of the peripheral immune compartment. A progressive decrease in cell cluster diversity was observed in association with disease severity: HD displayed greater immune cell diversity, whereas PBMCs from HTLV-1-infected patients, particularly those with acute ATL, are predominantly composed of T_h_ CD4+, indicating reduced heterogeneity (**Fig. 1a-b** and **Suppl. Fig. 2b**). T_h_ CD4+ cells dominated the landscape across all clinical groups but were particularly enriched in acute ATL (**Fig. 1b, 1c** and **Suppl. Fig. 2b**), suggesting clonal expansion and skewing of this compartment.

**Figure 1:**
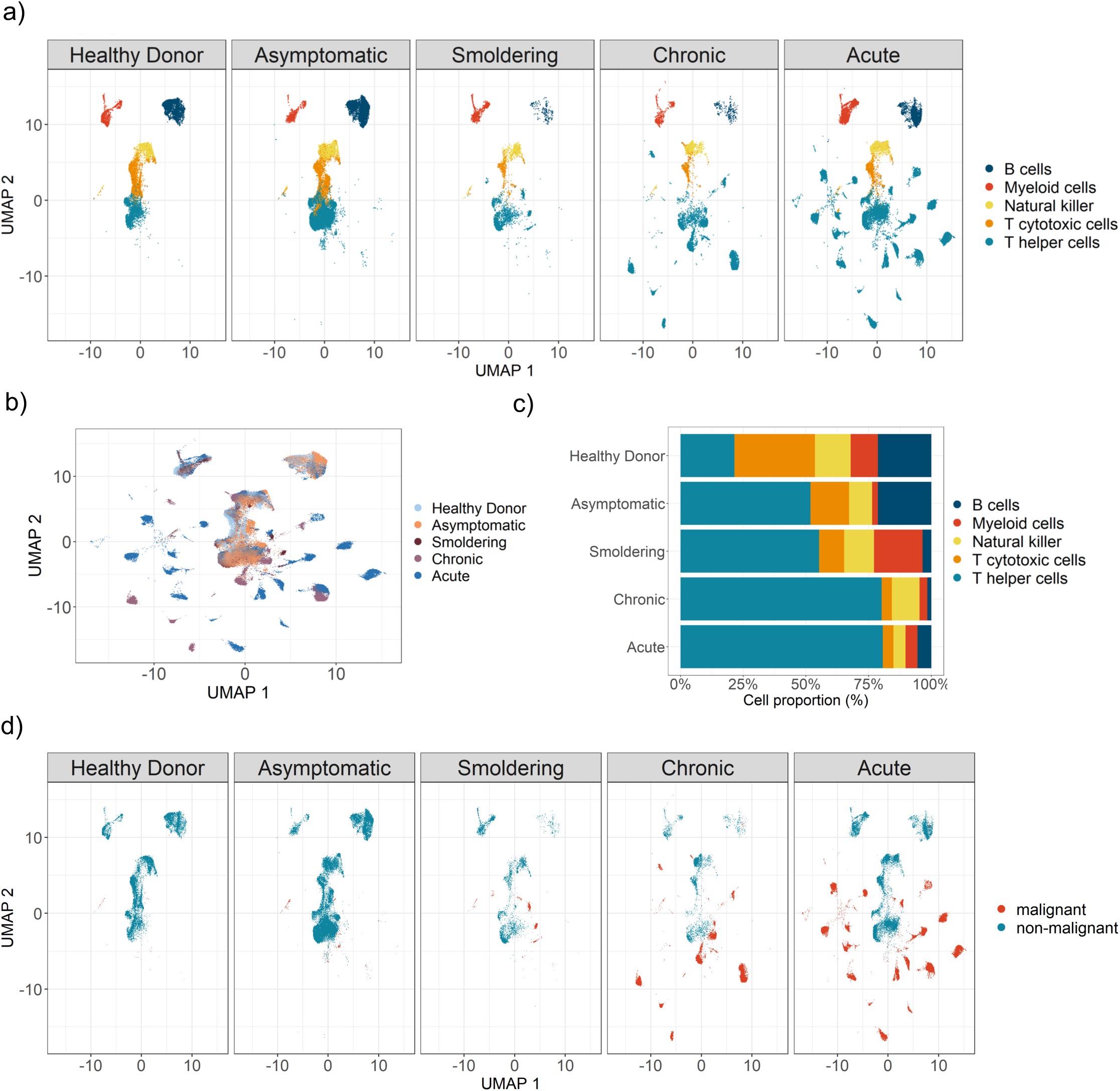
Single-cell analysis of PBMCs from HTLV-1 patients across disease stages. (a) UMAP visualization (reproduced from Koya et al., 2021) of PBMCs single-cell transcriptomes from healthy donors and HTLV-1 clinical subtypes (asymptomatic carriers, chronic ATL, and acute ATL), with cells colored by sample group. (b) UMAP colored by unsupervised clustering, showing immune cell clusters identified across all samples. (c) UMAP annotated by major immune cell types (e.g., CD4⁺ T helper, CD8⁺ T, NK, B cells, monocytes), based on marker-gene expression. Proportions of annotated cell types are shown for each clinical group. (d) UMAP highlighting malignant-like T-cell populations as defined in the original dataset, displayed by clinical subtype.

Chronic and acute ATL cases exhibit reduced immune heterogeneity due to the dominance of CD4⁺ T cells, in contrast to HD and ACs, who retain a more diverse PBMCs composition (**Fig. 1c**), indicating an immune architecture increasingly reshaped during disease progression. Concurrently, transcriptional features consistent with malignant transformation became more prominent along the disease trajectory. Cell clusters characterized by high expression of FOXP3, IL2RA, and PLCG1, markers classically associated with HTLV-1–driven leukemogenesis and ATL^1,31^, were enriched in patients with more advanced disease (**Fig. 1d** and **Suppl. Fig. 2c**).

### Peripheral Immune Cells Acquire Neuroimmune Transcriptional Identities Across HTLV-1 Disease Progression

Differential expression analysis revealed a progressive increase in the total number of differentially expressed genes (DEGs) across clinical stages of HTLV-1 infection, with the acute phase showing the greatest number of differentially expressed genes, particularly within T-cell subsets such as T_h_ CD4⁺. (**Fig. 2a**).

**Figure 2:**
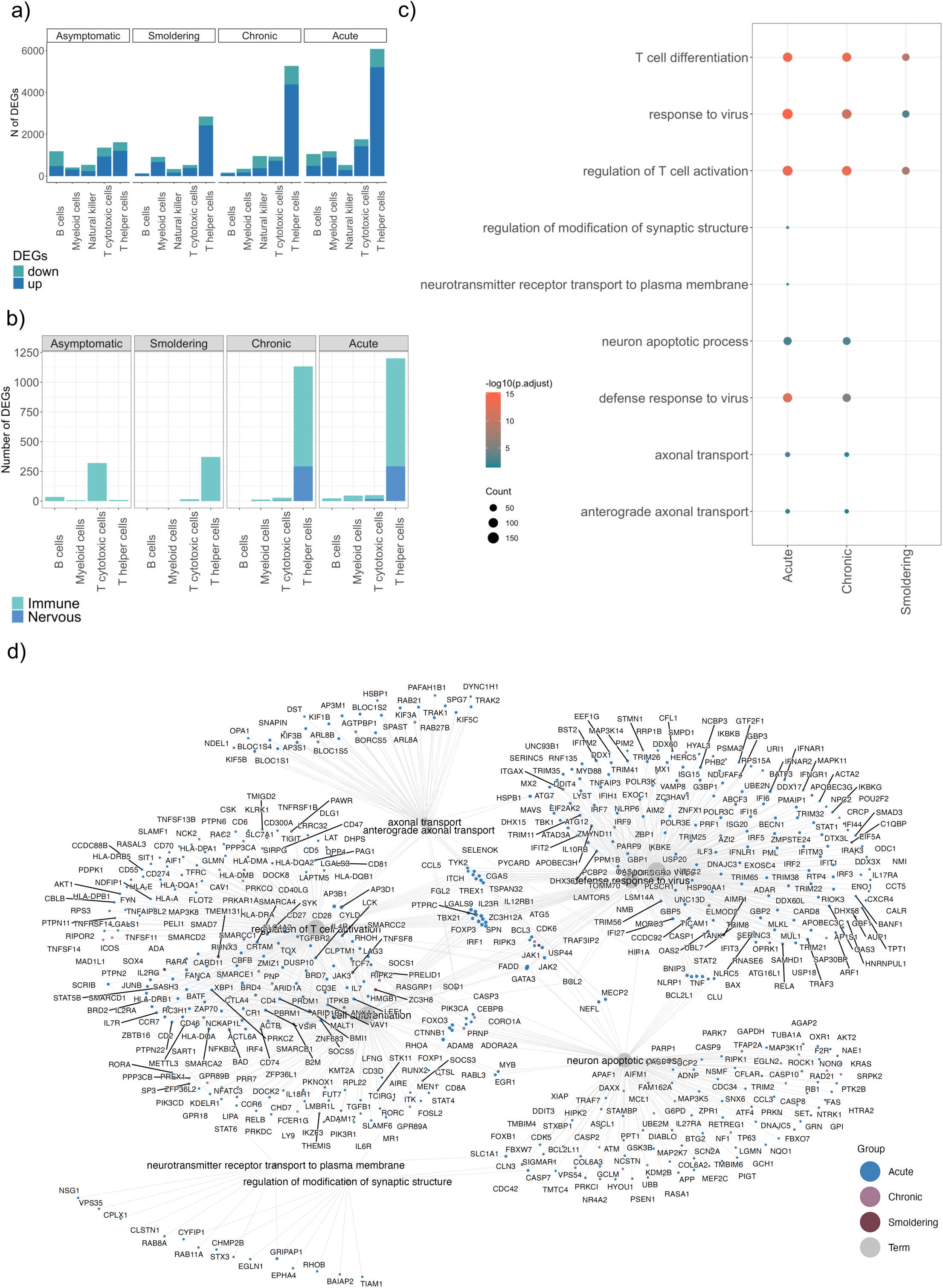
Neuro-immune genes and biological process signatures across HTLV-1 disease stages at the single-cell level. (a) Number of differentially expressed genes (DEGs) per immune cell subset in each clinical group compared to healthy donors. Bars indicate upregulated and downregulated DEGs in each comparison. (b) DEGs from each comparison categorized by functional annotation (immune-related, nervous system-related, or neuro-immune-related) and displayed by cell subset and clinical group. (c) Gene Ontology (GO) Biological Process enrichment analysis of DEGs for each immune cell subset and clinical group versus controls. Dot size indicates number of counts; color indicates -log10 adjusted p-value. (d) Network of DEGs and terms interaction from peripheral CD4⁺ T cells. Nodes represent genes; edges represent known or predicted associations derived from BPs database and co-expression.

The number of DEGs increased markedly across clinical stages, with the largest transcriptional remodeling occurring in the chronic and acute phases (**Fig. 2b**). Within this global shift, genes annotated to immune processes accounted for the vast majority of DEGs across all cell types and stages, suggesting that immune program remodeling is the dominant feature of HTLV-1 progression. In contrast, nervous system–associated DEGs comprised a smaller subset that became most evident in later disease stages, emerging predominantly within the T cell compartment, supporting a stage-dependent enrichment of neuroimmune-linked transcriptional signatures as disease severity increases (**Fig. 2b**).

Functional enrichment of these DEGs uncovered a robust neuroimmune program, with a substantial fraction of genes annotated to biological processes (BPs) typically confined to the central nervous system (**Fig. 2b**). Gene ontology enrichment analysis revealed that acute-phase T helper cells were characterized by significant high expression of biological processes associated with immune and nervous BPs (**Fig. 2c**).

These patterns contrasted with the other disease stages, as significant enrichment of neuroinflammatory biological processes was observed only in chronic cases, while ACs and smoldering stages showed no significant enrichment. Acute cases showed the strongest enrichment for nervous system-related terms, including synaptic structure, neurotransmitter transport, neuronal apoptosis, and axonal transport, whereas chronic cases displayed a more restricted neuronal signature. Both stages also shared immune-related terms, such as T cell differentiation, T cell activation, and antiviral responses. To explore the molecular programs contributing to neuroimmune dysregulation, we conducted a network-based analysis of DEGs to linkage between genes and terms (**Fig. 2d**).

To further illustrate this pattern, network analysis showed that the enriched terms in T helper cells were organized into interconnected immune and neuronal terms. Immune-related terms, including *response to virus*, *defense response to virus*, *response to type I interferon*, *T cell activation*, and *T cell differentiation*, formed a connected cluster, reflecting substantial overlap in antiviral- and activation-related genes such as *FOXP3, IL12RB1, JAK1, USP44,* and *GATA3*. Neuronal-associated terms, including *axonal transport*, *anterograde axonal transport*, *neuron apoptotic process*, *regulation of modification of synaptic structure*, and *neurotransmitter receptor transport to plasma membrane*, were likewise organized into a related subnetwork. Together, these connections indicate that antiviral and T-cell activation pathways converge with neuronal-associated processes through shared genes, including *BCL2, CYLD, SCRIB, CTNNB1, GATA3, FADD, ATG7, MYB,* and *EGR1*, supporting the existence of a coordinated neuroimmune transcriptional program.

### T Cells Adopt Synaptic Signaling During Neuroimmune Reprogramming in HTLV-1 Infection

The examples above underscore a tight coupling between canonical immune pathways and aberrantly activated neuron-like transcriptional programs in leukocytes. This suggest that neuroimmune crosstalk extends beyond bidirectional signaling between the nervous and immune systems, potentially being embedded within immune-cell regulatory states themselves. To further characterize the extent of neuroimmune reprogramming in HTLV-1 infection, we investigated whether peripheral immune cells adopt transcriptional programs commonly associated with neuronal synaptic signaling (**Supplementary Table 2**). Leveraging complementary enrichment frameworks, including ArchipelaGO, we systematically mapped the engagement of neurotransmission-related gene pathways across major immune subsets in individuals with HTLV-1 infection.

PBMCs DEGs were analyzed for pathway and functional enrichment to identify synapse-associated signatures linked to distinct HTLV-1 clinical outcomes (**Fig. 3a**). A bar plot summarizing the number of DEGs mapped to upregulated DEGs related to synaptic signaling pathways across immune cell types and HTLV-1 clinical subtypes is shown in **Figure 3b**. Overall, synaptic-pathway–associated DEGs were most prominent in T helper cells compartments, with comparatively fewer genes detected in B cells, T cytotoxic cells and NK cells across ATL groups (**Fig. 3b**). Among all groups, the Acute subtype displayed the greatest accumulation of synapse-related DEGs, most prominently in T helper cells, with substantial representation of cholinergic, dopaminergic, and GABAergic gene signatures. In contrast, DEG counts were generally low in the Asymptomatic and Smoldering groups, while the Chronic group displayed an intermediate profile characterized mainly by higher counts in T helper cells, remaining below the Acute group overall. (**Fig. 3b and Supplementary Table 2**).

**Figure 3:**
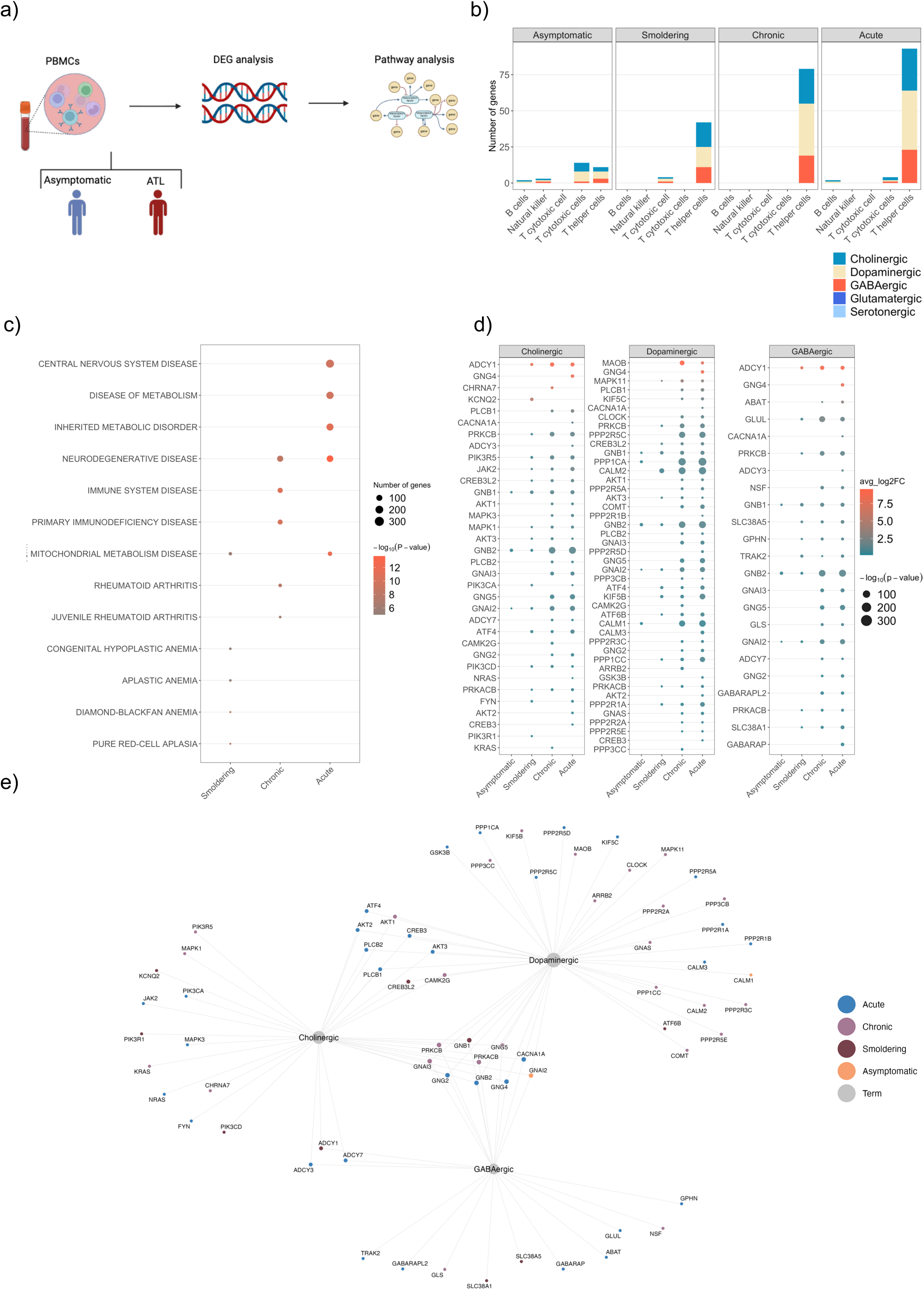
Enrichment analysis of synaptic and neuroimmune pathways in HTLV-1–associated T helper cells. (a) DEGs were subsequently analyzed using pathway/functional enrichment to characterize coordinated molecular programs distinguishing asymptomatic carriers, ATL, and HAM/TSP groups. (b) Stacked bars represent the number of DEGs per immune cell population within each clinical group (Asymptomatic, Smoldering, Chronic, Acute). Colors denote the synaptic pathway annotation of each DEG (cholinergic, dopaminergic, GABAergic, glutamatergic, serotonergic). The plot highlights stage- and cell-type–dependent differences in the contribution of synaptic signaling gene sets. (c) Bubble plot of disease-enrichment results based on the Jensen DISEASES Curated 2025 gene-set library across HTLV-1 clinical groups (Asymptomatic, Smoldering, Chronic, and Acute) using the total set of DEGs. Each dot represents one enriched disease term within a given group; dot size indicates the number of associated genes, and color represents enrichment significance. (d) Bubble plot of upregulated genes associated with synaptic pathways across HTLV-1 clinical groups (Asymptomatic, Smoldering, Chronic, and Acute). Each dot represents one gene within a given pathway and group; dot size indicates enrichment significance (−log10(P-value), and color represents the average log2 fold change. The Acute group showed the broadest and strongest transcriptional activation, particularly in the Dopaminergic and GABAergic pathways, whereas the Chronic group displayed an intermediate pattern and Smoldering and Asymptomatic groups showed fewer significant changes. Notably, genes such as *ADCY1, GNG4, MAOB, MAPK11, GLS, GLUL*, and *ABAT* contributed prominently to the acute synaptic signature (e) Network representation of synapse-related genes associated with the Cholinergic, Dopaminergic, and GABAergic pathways. Gray nodes represent pathway terms and connected gene nodes indicate pathway-associated genes. Node colors denote disease groups (Acute, Chronic, Smoldering, and Asymptomatic). The network highlights a predominance of Acute-associated genes, particularly in the Dopaminergic and GABAergic modules, while Chronic and Smoldering genes show more limited contributions and Asymptomatic genes are sparse. Shared genes across pathways suggest coordinated regulation of synaptic signaling.

Analysis using Jensen DISEASES Curated 2025^32^ revealed that the Acute group exhibited the strongest disease enrichment, especially for neurological and metabolic terms, while the Chronic group showed an intermediate profile, the Smoldering group displayed weak enrichment, and the Asymptomatic group showed no significant disease enrichment (**Fig. 3c**).

Analysis restricted to upregulated genes revealed a progressive increase in synapse-related transcriptional activity across disease groups, with the strongest signal observed in the Acute subtype. This group showed the largest number of significant genes and the highest overall expression changes, especially in the dopaminergic and GABAergic pathways. The Chronic group displayed an intermediate pattern, whereas the Smoldering and Asymptomatic groups showed only limited or minimal enrichment. Notably, several genes involved in synaptic signaling, including ADCY1, GNG4, MAOB, CHRNA7, and GLUL, were among the most prominently upregulated, supporting enhanced neurotransmission-related activity in the Acute group (**Fig. 3d**).

To resolve these transcriptional features within PBMCs clusters, ArchipelaGO-based enrichment mapping on UMAP cell clusters was performed on acute ATL samples. This analysis mapped synapse-related biological processes, including synaptic transmission, neurotransmitter transport, neuropeptide signaling, synapse organization, and peripheral nervous system development, to discrete cellular clusters enriched in malignant T cells (**Suppl. Fig. 3a**). These processes were not associated with the main non-malignant immune clusters in the same analysis. To enhance our analysis, we subset the T cell clusters to assess differences in synaptic-related pathways across clinical conditions. UMAP projections illustrate the overall cellular landscape: when cells are colored by clinical group (HD, Asymptomatic, Smoldering, Chronic, and Acute), we observe partial overlap between groups alongside condition-specific enrichment in distinct regions of the embedding (**Suppl. Fig. 3b**).

Notably, acute and chronic samples show broader dispersion, suggesting increased transcriptional heterogeneity compared with HD and AC samples. When cells are colored by T cell subtype, T_h_ CD4+ and T_c_ CD8+ cells segregate into clearly distinct regions, indicating that cell identity is a major driver of transcriptional variation within the T cell compartment (**Suppl. Fig. 3c**).

Together, these results suggest a progressive increase in neuronal-related gene expression from HD and asymptomatic states toward chronic and acute conditions. This pattern indicates that disease severity is associated with enhanced activation of neuronal signaling-related transcriptional programs, particularly in more advanced clinical stages.

The network analysis revealed three major hubs corresponding to the Cholinergic, Dopaminergic, and GABAergic pathways, with most connected genes belonging to the Acute group and a smaller contribution from the Chronic group. In the cholinergic module, genes involved in intracellular signaling and synaptic regulation, including *PIK3CA, PIK3CD, PIK3R1, PIK3R5, MAPK1, MAPK3, JAK2, FYN, CHRNA7,* and *ADCY1/ADCY3/ADCY7*, were prominently represented (**Fig. 3e**). The dopaminergic module appeared as the most expanded network, including genes such as *MAOB, COMT, GNAS, CALM1/2/3, CLOCK, MAPK11, GSK3B, KIF5B,* and *KIF5C*, together with several phosphatase-related genes from the *PPP2R* and *PPP3* families, suggesting broad regulation of dopamine-related signaling (**Fig. 3e**). In the GABAergic module, genes associated with inhibitory synaptic transmission and neurotransmitter metabolism, such as *GPHN, NSF, ABAT, GLS, GLUL, GABARAP, GABARAPL2, SLC38A1, SLC38A5,* and *CACNA1A*, were identified (**Fig. 3e**). Several signaling genes, including *PLCB1, PLCB2, AKT1, AKT2, AKT3, CREB3, CREB3L2,* and *CAMK2G*, were shared across pathways, indicating coordinated synapse-related regulation. Overall, these results suggest that synaptic dysfunction is mainly driven by genes from the Acute group.

### ATL Displays a Dominant Neuroimmune Transcriptomic Signature Revealed by Integrative Meta-analysis

To extend the neuroimmune patterns observed in single-cell analyses, we conducted a comprehensive meta-analysis integrating publicly available transcriptomic datasets from PBMCs of individuals with HTLV-1 infection. Using a PRISMA-guided selection process (**Suppl. Fig. 4**), we screened 536 datasets and curated a final panel of six, which included both bulk RNA-seq and MicroArray datasets from ACs, patients with ATL, and individuals diagnosed with HAM/TSP. The total number of individuals across these datasets is as follows: 100 from EGAS00001004936, 45 from GSE29312, 20 from GSE38537, 16 from GSE233437, and 50 from GSE33615. The subtype distribution includes 41 individuals with Acute, 46 with Chronic, 6 with Smoldering, 27 Asymptomatic, 38 HD, 6 with Lymphoma, and 18 with HAM/TSP. Datasets were harmonized using ExpressAnalysis, with batch effect corrections enabling high-confidence cross-condition comparisons (**Suppl. Fig. 5a, 5b and Supplementary Table 1 and 3**).

Differential expression analysis across cohorts revealed that ATL exhibits the most substantial transcriptomic shift, with 5,884 genes downregulated and 2,138 upregulated, compared to 551 downregulated and 2,430 upregulated in asymptomatic individuals, and 849 downregulated and 1,383 upregulated in HAM/TSP samples (**Suppl. Fig. 5c**). We quantified DEGs related to immune, nervous, and overlapping neuroimmune processes in individuals with HTLV-1-associated conditions using BPs ontology.

**Figure 4a** summarizes the workflow used for the functional enrichment analyses shown in **Figure 4**. Consistent with the DEG counts, ATL displayed the largest transcriptional perturbation, with the highest numbers of both immune- and nervous system–annotated DEGs (**Fig. 4b**). Asymptomatic carriers showed an intermediate DEG burden with a comparatively small nervous-system component, whereas HAM/TSP exhibited the lowest overall DEG numbers while still retaining a detectable nervous-system–associated signal (**Fig. 4b**).

**Figure 4:**
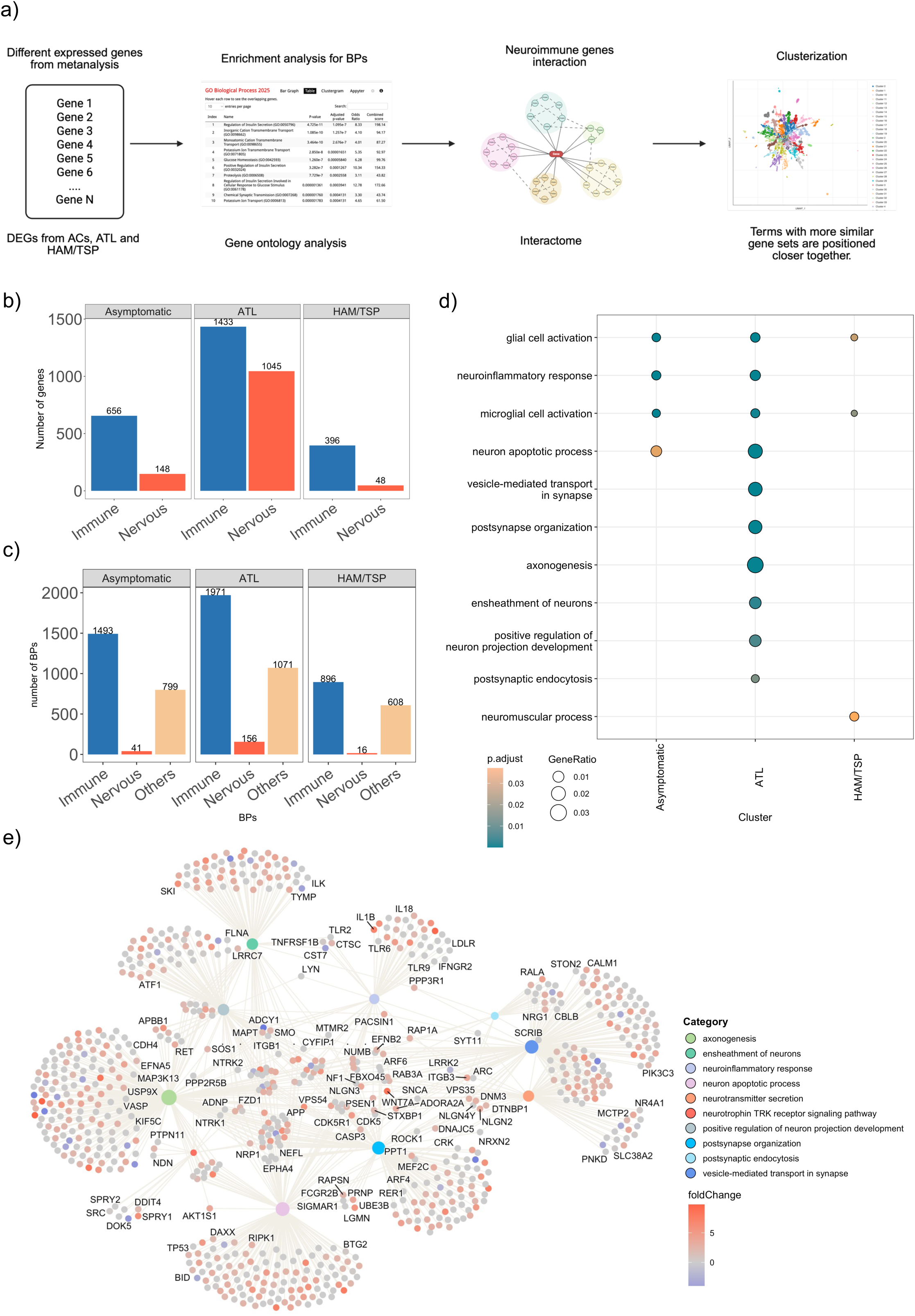
Meta-analysis of transcriptomic datasets and neuroimmune functional annotation across HTLV-1 clinical groups. (a) A gene list of DEGs was submitted to Gene Ontology Biological Process (GO-BP) enrichment analysis, resulting in table of significantly enriched terms. These enriched terms are then organized into a network (interactome) based on gene overlap, revealing coherent functional modules. Finally, this structure is projected onto a two-dimensional map to summarize and compare the main biological programs represented by the input gene set. (b) The number of DEGs per clinical comparison versus controls is grouped by immune-related, nervous system-related, or neuroimmune-related annotations. (c) GO enrichment of DEGs from each clinical group compared to healthy donors, highlighting nervous system-related biological processes. (d) GO Biological Process terms related to neuroimmune functions that are enriched in each clinical comparison versus healthy donors, along with a cumulative number of significantly enriched GO Biological Process terms per functional class across clinical groups. (e) A functional interaction network of enriched GO terms and associated genes across PBMCs datasets, grouped by annotation category.

Although our single-cell analyses indicate that the most substantial transcriptional changes are concentrated in T-cell populations, the PBMCs-level meta-analysis remains significant even when additional immune compartments are included. This robustness likely reflects (i) the high abundance of T cells within PBMCs, which strongly influences bulk signals, and (ii) the magnitude and consistency of ATL-associated neuroimmune transcriptional programs across datasets, allowing these signatures to remain detectable at the whole-PBMCs level despite cellular heterogeneity.

Consistently, functional categorization of DEGs shows that ATL patients display the highest number of BP terms associated with immune, nervous, and neuroimmune processes, suggesting broad transcriptional activation across these systems. In contrast, asymptomatic carriers exhibit a more balanced profile, and HAM/TSP patients present fewer BPs overall, with limited nervous involvement (**Fig. 4c**). Together, these findings align with and further support our neuroimmunoma hypothesis.

Meanwhile, **Figure 4d** shows that ATL harbors the most pronounced enrichment and the largest shifts in nervous system–related biological processes, spanning multiple neuronal/synaptic terms, whereas ACs and HAM/TSP exhibit only secondary, more modest changes in these nervous-associated programs. Notably, inflammatory pathways, including microglial-associated signatures such as microglial cell activation and neuroinflammatory response, display a distinct distribution across groups (**Fig. 4d and Supplementary Table 4**).

To interrogate neuroimmune transcriptional programs in ATL, we constructed a functional enrichment interactome from PBMCs-derived DEGs, linking significantly enriched terms through shared genes (gene nodes shaded by fold-change). The network resolved into a modular, biologically coherent architecture, featuring (i) a prominent neuronal/synaptic module encompassing postsynapse organization, and endocytosis, vesicle-mediated transport in synapses, neurotransmitter secretion, and neurotrophin–TRK signaling; (ii) a neuroinflammatory module; and (iii) an apoptosis-associated module. Additional connectivity around axonogenesis/neurite outgrowth and neuron ensheathment terms further supports the coordinated dysregulation of neuroglial-related pathways in peripheral immune cells (**Fig. 4e and Supplementary Table 4**).

Taken together, these results reconcile single-cell and PBMCs-level observations by indicating that, while T cells exhibit the strongest cell-intrinsic changes, the ATL-associated neuroimmune program is sufficiently robust and partially shared across PBMCs compartments to remain detectable in integrated whole-PBMCs transcriptomes despite cellular heterogeneity. Notably, this signal was consistently recovered across platforms, as both bulk RNA-seq and microarray-based meta-analyses captured similar ATL-associated transcriptional shifts, further underscoring the robustness of the dominant systemic neuroimmune signature in ATL.

### Neuroimmune Genes Discriminate ATL and Reflect Functional Crosstalk Between Leukemia and Nervous System Programs

To assess the potential involvement of neuroimmune signaling in HTLV-1–associated leukemogenesis, we performed comprehensive statistical and machine learning analyses on peripheral blood transcriptomes from patients with acute and chronic ATL using dataset GSE33615. Beginning with a curated panel of 1,382 neuroimmune-related metaDEGs (**Supplementary Table 5**), we performed PCA and applied the Kaiser criterion (eigenvalues > 1), identifying two principal dimensions (Dim1 and Dim2) that separated ATL samples from healthy donors. This separation was driven by the highest-loading features on these axes, with significant contributions from genes including *TIAM2*, *ZNF365*, *PPP1BP1*, *SKIL*, *AIM2*, SPRY1, *BTG2*, *CXCR4*, *SSNA1*, *ATF4*, *XRCC5*, *SETX*, *ATP6V0D1*, and *PRKCH* (**Fig. 5a–b**).

**Figure 5:**
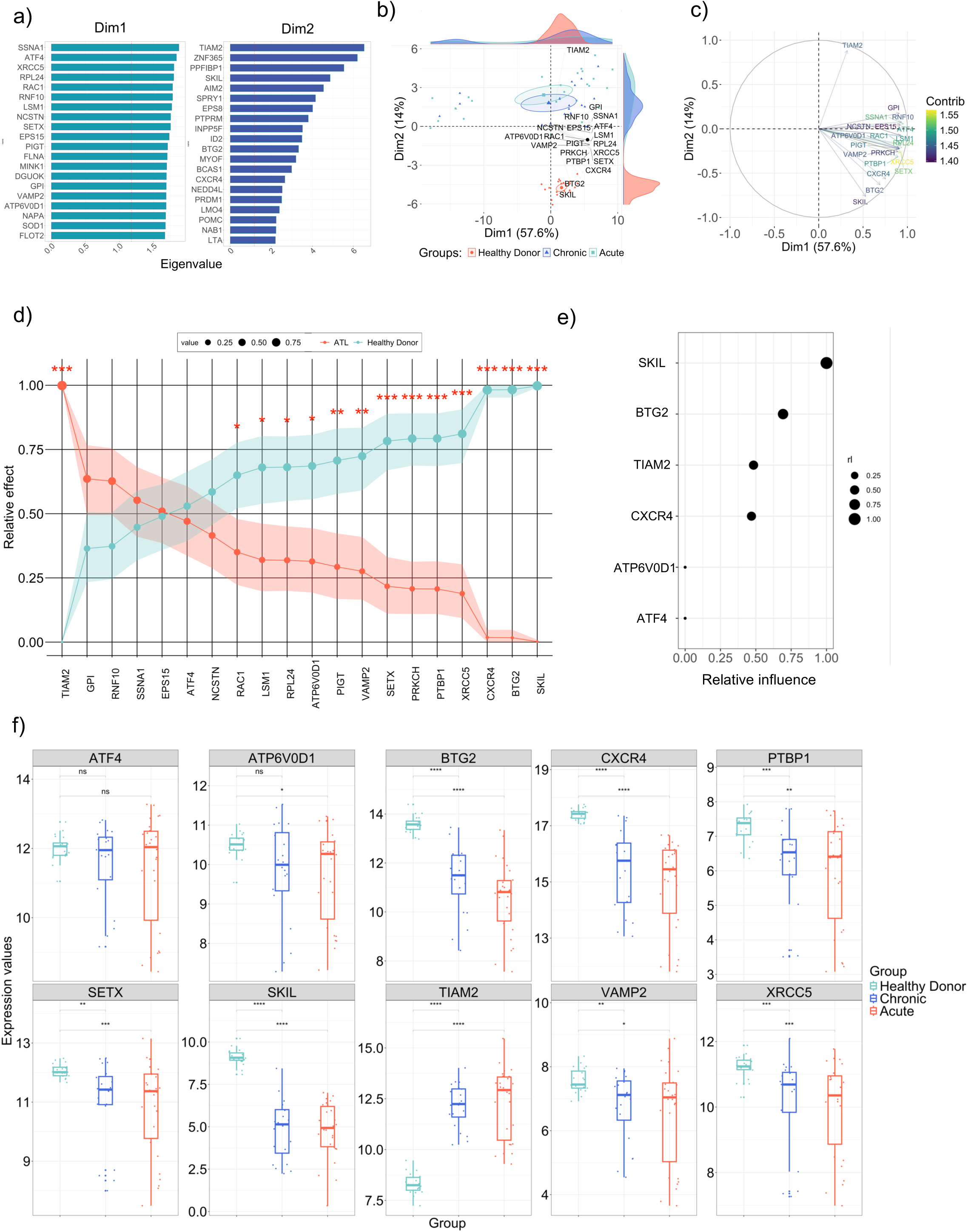
Integrated machine learning, and statistical modeling identify neuroimmune-related intracellular signatures in HTLV-1 infection. (a) Principal Component Analysis (PCA) of neuroimmune gene expression across controls and ATL samples, highlighting genes that contribute most to Dim1 and Dim2. (b) The top 20 neuroimmune genes selected based on their eigenvalue contribution to PCA group separation. (c) A PCA loading plot displaying the direction and magnitude of gene contributions to Dim1 and Dim2; the color scale indicates contribution (“Contrib”). (d) Relative effect analysis for PCA-selected neuroimmune genes comparing ATL and control groups; circle size indicates the relative effect magnitude. (e) Gradient Boosting Machine (GBM) model illustrating the relative influence of neuroimmune genes for group classification. (f) Boxplots of expression levels of selected neuroimmune genes across acute ATL, chronic ATL, and healthy donors. Statistical comparisons were performed using Wilcoxon rank-sum tests; p < 0.05 (\**), p < 0.01 (*\*\**), p < 0.001 (*\*\*\*), and p < 0.0001 (****), and “ns” for non-significant.

These genes were prioritized because they (i) showed consistent differential expression in ATL across datasets and (ii) are functionally linked to pathways relevant to HTLV-1 pathogenesis, including T-cell signaling and differentiation (*TIAM2, CXCR4*), stress and integrated stress-response programs (*ATF4, BTG2, ATP6V0D1*), genome maintenance (*XRCC5, SETX*), and transcriptional regulation of immune activation and TGF-β–associated responses (*SKIL*).

In **Figure 5c**, each arrow represents the loading vector of a gene, reflecting both the direction and strength of its contribution to the variance captured in the reduced dimensional space. Genes such as *TIAM2, SKIL, CXCR4, XRCC5, BTG2, SETX* and *PTBP1* exhibit pronounced loadings along Dim1 and/or Dim2, indicating their significant role in driving the transcriptomic stratification between ATL patients and HD (**Fig. 5c**). The color scale denotes each gene’s relative contribution to the principal components, with warmer hues (yellow to green) corresponding to higher contributions.

To evaluate the discriminatory power of these candidate genes, we performed a bootstrapped multivariate analysis of variance (MANOVA), which identified *TIAM2, CXCR4, BTG2, and SKIL* as genes with robust and statistically non-overlapping effects between ATL and control samples (**Fig. 5d and Supplementary Table 6**). These findings were further supported by gradient boosting machine (GBM) modeling, which ranked the relative influence of genes on group classification. *TIAM2, BTG2, CXCR4*, and *SKIL* were consistently identified as the most influential features, along with *ATP6V0D1* and *ATF4* (**Fig. 5e**).

Notably, *SKIL* (Ski-like proto-oncogene) is a transcriptional coregulator with established links to TGF-β/SMAD signaling and regulation of proliferation and immune-cell differentiation^33^. Given the central role of TGF-β–associated programs in T-cell homeostasis and leukemic reprogramming, the strong contribution of *SKIL* in both MANOVA and GBM suggests that disruption of this regulatory axis may be an important component of ATL-associated neuroimmune remodeling, beyond its statistical separation power alone.

In addition, statistical comparisons using the Wilcoxon rank-sum test revealed significant differential expression in nine out of the ten genes analyzed across clinical groups. Specifically, eight genes showed statistically significant differences when comparing both acute and chronic ATL groups to HD (**Fig. 5f and Supplementary Table 6**). Notably, *ATF4* was the only gene that did not reach statistical significance in either comparison (**Fig. 5f**).

Peripheral blood immune cells from HD, AC, and ATL were previously profiled by single-cell RNA-seq and clustered by lineage to establish the major immune compartments and enable condition-wise comparisons (**Fig. 6a**). Here, we build directly on that framework by restricting the analysis to a focused set of twelve neuroimmune genes and quantifying how their expression shifts across conditions within each lineage-defined subset (**Fig. 6b**). Using the group means (central bars), ATL showed the most consistent repression in T helper cells, where SETX, SKIL, XRCC5, ATF4, ATP6V0D1, BTG2, and PTBP1 were significantly reduced compared with ACs and/or HDs (**Fig. 6b**), suggesting a selective loss of multiple regulatory programs in ATL-associated CD4⁺ T cells.

**Figure 6:**
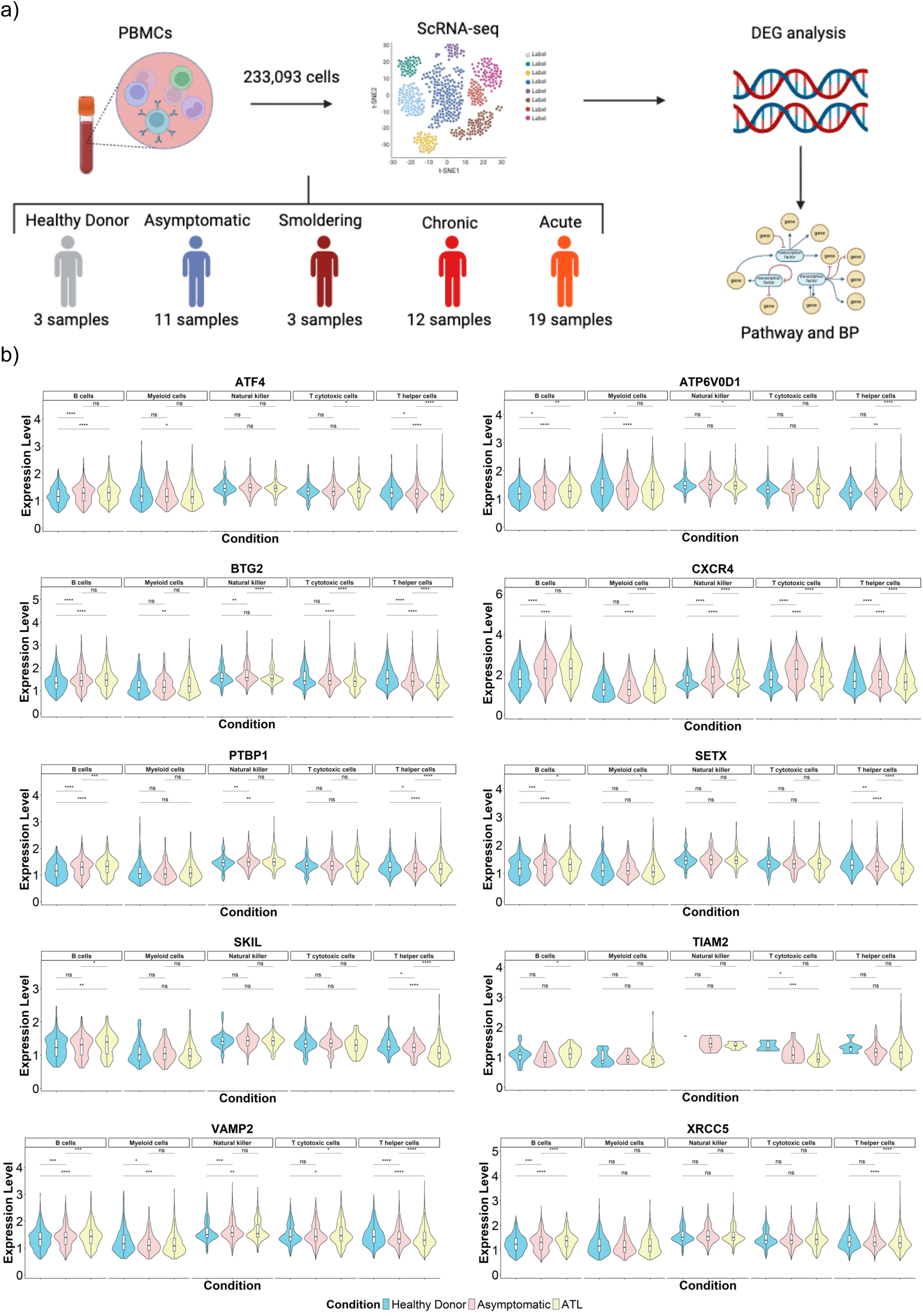
Cell-Type-Specific Gene Expression Patterns Reveal Immune Signatures Across HTLV-1 Conditions: (a) Analytical workflow of the study. (b) Violin plots depict the expression profiles of twelve genes (ATF4, ATP6V0D1, BTG2, CXCR4, PTBP1, SETX, SKIL, TIAM2, VAMP2, and XRCC5) across five major immune cell populations: B cells, myeloid cells, natural killer (NK) cells, T cytotoxic cells, and T helper cells, under three clinical conditions: Asymptomatic (red), Healthy donors (blue), and Tumor (yellow). The overall distribution of gene expression across all cell types is represented by the black violins. Statistical comparisons between conditions were performed using the Wilcoxon rank-sum test, with significance levels indicated as follows: p < 0.05 (), p < 0.01 (), p < 0.001 (), p < 0.0001 (****), and non-significant (ns).

In contrast to these repressed genes, CXCR4 was not reduced in ATL T helper cells; instead, its mean expression was maintained and, in several compartments, increased (**Fig. 6b**). Specifically, CXCR4 was significantly upregulated in myeloid cells, NK cells, and cytotoxic T cells in ATL relative to HDs, consistent with heightened chemokine-axis activity across non–CD4 compartments (**Fig. 6b**). PTBP1 showed cell-type–restricted behavior, with reduced expression in ATL T helper cells but higher expression in B cells and NK cells, supporting divergent post-transcriptional regulation across peripheral immune niches (**Fig. 6b**).

Additional gene-level patterns reinforced this compartment specificity. SETX, SKIL, and XRCC5 were not significantly reduced in cytotoxic T cells, despite clear changes in T helper cells (**Fig. 6b**). TIAM2 was not elevated in T cell populations; it was significantly downregulated in cytotoxic T cells while being significantly increased in B cells from ATL compared with ACs (**Fig. 6b**). VAMP2 showed more modest, context-dependent differences without a consistent ATL-wide repression pattern across all subsets (**Fig. 6b**).

### Flow Cytometric Profiling and Correlation Analysis of Neuroimmune Markers in PBMCs Reveals Distinct Signatures Across HTLV-1 Clinical Outcomes

To test the neuroimmune-related signature identified in our primary analysis, we performed an independent cross-cohort evaluation using PBMCs from a UK cohort and validated key transcript-level signals at the protein level by multiparameter flow cytometry in PBMCs from HD, AC, ATL, and HAM/TSP (**Supplementary Table 7**). These proteins were selected based on their loading values in PCA, biological relevance in immune–neural crosstalk, and predictive power in the GBM model (**Fig. 6**).

At the protein level, V-ATPase D1 exhibited one of the most consistent alterations, showing a marked reduction in HTLV-1–infected groups relative to HD (**Fig. 7a**), along with decreased signal intensity across infected samples (**Fig. 7b**). ATF4 displayed a stage-dependent pattern, with higher levels in ACs but lower levels in ATL (**Fig. 7a–b**). SKIL was profoundly suppressed across HTLV-1–infected groups (near-minimal detection compared to HD; **Fig. 7a–b**). CXCR4, PTBP1, SETX (Senataxin), and XRCC5/Ku80 also demonstrated overall reductions in HTLV-1 infection (**Fig. 7a–b**), with SETX showing a modest relative increase in ATL compared to AC and HAM/TSP that approached significance (**Fig. 7a**). In contrast, VAMP2 and Ku80 remained broadly expressed across groups (>95% positive), but both exhibited significantly reduced gMFI in HTLV-1–infected individuals compared to HD, indicating preserved marker frequency but decreased per-cell signal intensity (**Fig. 7b**).

**Figure 7:**
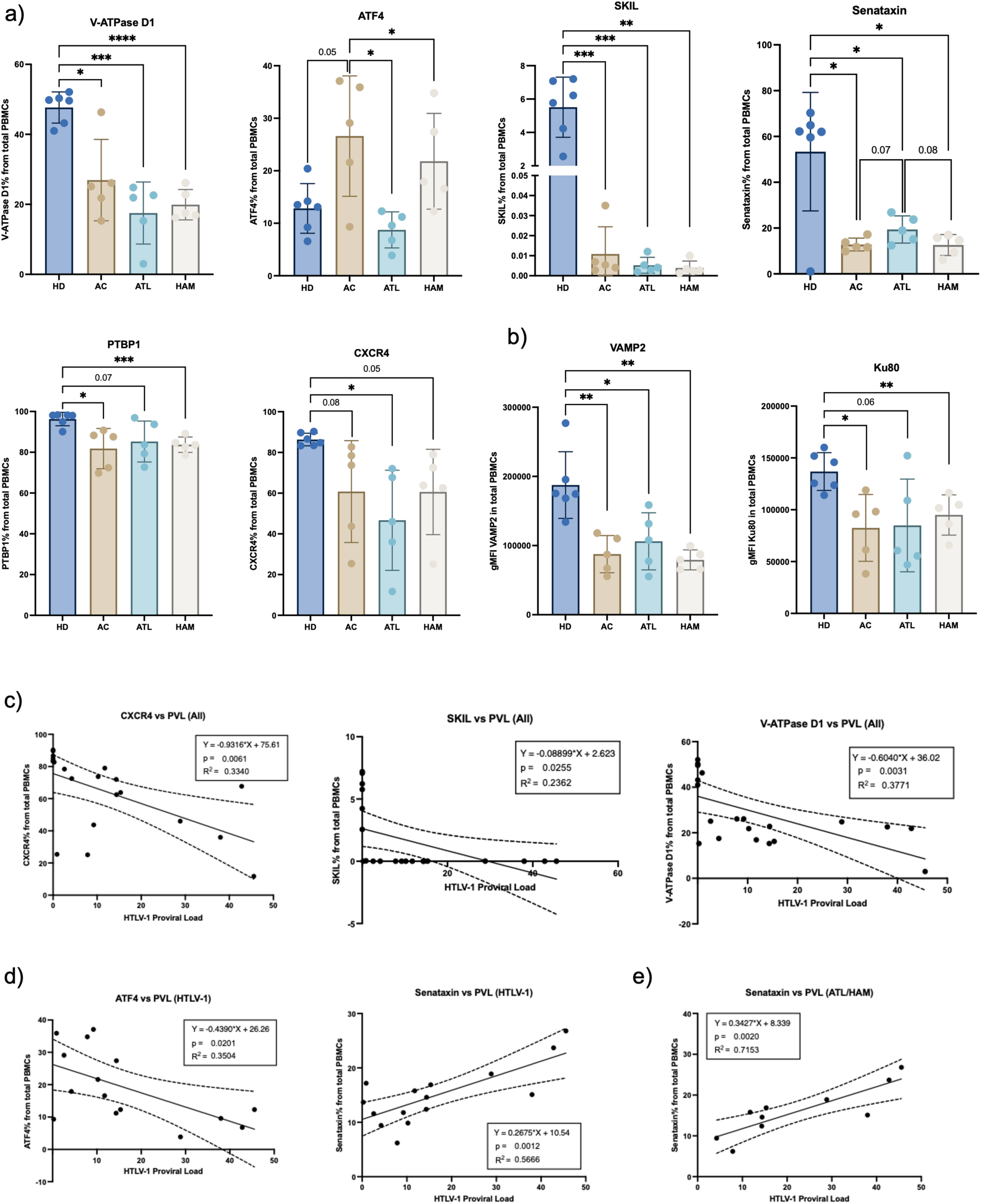
Flow cytometry quantification of neuroimmune and stress-response markers in PBMCs from HTLV-1–infected individuals and association with proviral load. Multiparameter flow cytometry was employed to quantify intracellular neuroimmune and stress-response markers in PBMCs from healthy donors (HD), asymptomatic carriers (AC), HAM/TSP, and ATL individuals. (a) Frequencies of PBMCs positive for ATP6V0D1, ATF4, SKIL, SETX, PTBP1, and CXCR4 across clinical groups. (b) Geometric mean fluorescence intensity (gMFI) of intracellular VAMP2 and XRCC5 (Ku80) across groups. Points represent individual donors; bars show mean ± SD. Group comparisons were conducted using unpaired t-tests with Welch’s correction; p < 0.05 (\**), p < 0.01 (*\*\**), p < 0.001 (*\*\*\*), and p < 0.0001 (****). (c–e) Linear regression analyses between HTLV-1 proviral load and marker expression are shown for: (c) all individuals, (d) HTLV-1–infected individuals only, and (e) symptomatic HTLV-1 infection (HAM/ATL). Each point represents one subject, and regression lines with 95% confidence intervals are displayed.^68^

Finally, proviral load (PVL) correlated with several of these protein changes, linking viral burden to disease severity: in longitudinal follow-up, all individuals who later developed HAM/TSP already had high PVL (>1%) before symptom onset, supporting PVL as a risk marker for progression^33^. Additionally, elevated PVL has been associated with leukemic risk, where a “high-risk” HTLV-1 carrier subgroup with higher PVL showed a markedly increased long-term incidence of ATL, indicating that PVL rises in the clinical context most prone to malignant transformation^34^. Across all participants, increasing PVL was associated with a lower frequency of CXCR4⁺, SKIL⁺, and V-ATPase D1⁺ PBMCs (**Fig. 7c**). When restricting the analysis to HTLV-1–infected individuals, ATF4⁺ cell frequency negatively correlated with PVL, whereas Senataxin⁺ cell frequency showed a strong positive correlation with PVL (**Fig. 7d**). This Senataxin–PVL relationship remained evident in symptomatic cases (ATL/HAM), supporting Senataxin as a dynamic marker that scales with higher viral burden in clinically overt disease (**Fig. 7e**).

## Discussion

These findings indicate that HTLV-1 infection induces transcriptional alterations in genes and pathways associated with the neuroimmune response in peripheral immune cells, with these signatures becoming progressively more pronounced across the clinical spectrum from asymptomatic carriers to ATL. This dysregulated gene expression network, termed the neuroimmunome, converges on signaling pathways typically restricted to the nervous system, such as synaptic transmission, neuropeptide signaling, and neurotransmitter transport.

Our study builds on previous evidence of neuroimmune crosstalk in chronic viral infections and cancers ^35,36^, but extends this understanding by applying a systems biology lens to reveal how these interactions evolve across clinical stages. Through single-cell transcriptomics, we observed that CD4⁺ T_h_ cells, CD8⁺ T_c_ cells, and myeloid cells in ATL acute patients exhibit enriched expression of genes typically involved in dopaminergic, glutamatergic, serotonergic, and GABAergic synapses. These data indicate that malignant CD4⁺ T cells exhibit expression of genes involved in neurotransmission; however, our findings remain hypothesis-generating and require independent validation. These findings indicate that peripheral blood leukocytes express genes linked to biological processes involved in nervous system responses. Similar neuroimmune reprogramming has been described in other neuroinflammatory diseases, such as multiple sclerosis and Alzheimer’s disease, in which immune cells acquire neural-like characteristics that may contribute to both inflammation and neuronal dysfunction.^37,38^.

Our integrative analysis reveals that several neuroimmune genes, namely *ATF4*, *ATP6V0D1*, *SKIL*, *SETX*, *CXCR4*, and *VAMP2,* display both transcriptional and translational dysregulation in PBMCs of individuals with HTLV-1 infection, mapping a molecular trajectory that connects oncogenesis with neurological deterioration. Of particular interest, ATF4, a key effector of the integrated stress response, was elevated in ACs compared to HD. Although ATF4 levels also trended lower in ATL patients and higher in HAM/TSP patients relative to HD, these differences were not statistically significant.

Conversely, there was a significant reduction of ATF4 expression in both ATL and HAM/TSP patients when compared to ACs. This biphasic expression mirrors findings in Alzheimer’s disease, where ATF4 has been detected in axons and linked to the propagation of neurodegenerative pathology^39,40^. In the context of HTLV-1 infection, this pattern may signify early neuroprotective or immune-adaptive responses that collapse under chronic viral pressure, thereby facilitating both malignant transformation and neuroinflammatory sequelae.

The observed reduction of V-ATPase D1, a core subunit of the vacuolar ATPase complex, across all HTLV-1 infection groups further underscores the convergence of viral oncogenesis with neurodegenerative mechanisms. Dysfunctions in V-ATPase are known to disrupt lysosomal acidification and neuronal homeostasis, contributing to a wide spectrum of brain disorders, including epilepsy and encephalopathies^41,42^. In our study, V-ATPase D1 (ATP6V0D1) expression was most profoundly reduced in ATL and HAM/TSP, potentially reflecting impaired vesicular trafficking and lysosomal/endosomal acidification that are essential for immune-cell metabolism, antigen processing, and signaling^43,44^. Given that V-ATPases regulate endo-lysosomal function and vesicle dynamics across cell types, their dysfunction is linked to reduced cellular stress tolerance and proteostasis, and in neurons specifically, to defective synaptic vesicle loading and neurodegenerative phenotypes^43–45^.

Furthermore, *SETX* is an RNA/DNA helicase that resolves R-loops and limits transcription-associated genome instability^46^, whereas *XRCC5* is a core *NHEJ* factor essential for repairing DNA double-strand breaks^47^. *SKIL* is a SMAD-interacting transcriptional coregulator within the TGF-β/SMAD pathway, shaping transcriptional programs linked to cell-state control and genome maintenance^48^, while *PTBP1* is a major regulator of pre-mRNA splicing and RNA metabolism^49^. Collectively, these coordinated decreases are consistent with altered RNA-processing and DNA-repair/homeostasis pathways in HTLV-1–associated conditions.

In conjunction with extensive evidence that HTLV-1 proteins Tax and HBZ reprogram host immune signaling and sustain chronic inflammatory states, which drive both systemic immune dysfunction and CNS damage in HAM/TSP, these findings support a model in which HTLV-1 preferentially disrupts key neural-immune interface pathways propagating immune dysregulation and neural impairment^45^.

Beyond stress response and vesicular regulation, genes such as *CXCR4,* and *SETX* emerged as critical players in the immunoneural landscape of HTLV-1 infection. CXCR4, despite being reduced in all HTLV-1 infection groups compared to HD, was slightly elevated in ACs and HAM/TSP compared to ATL patients. *CXCR4* is essential for chemokine signaling and leukocyte migration and is implicated in microglial dysregulation across neurodegenerative diseases^50^.

Its role in ATL cell homing and in fostering neuroinflammation within the CNS aligns with our data showing inverse correlations between CXCR4 expression and HTLV-1 PVL, suggesting that its downregulation may indicate a loss of immune surveillance and neuroprotective function^51,52^. Similarly, Senataxin, a protein involved in resolving transcriptional stress and previously linked to neurological disorders like ataxia and neuropathies^53^, demonstrated a significant decrease in expression across all HTLV-1 infection groups compared to HD.

Although our study is systemic and exploratory and therefore does not establish causality, the altered PBMCs programs we observe are compatible with several well-described pathways through which peripheral immune activity can relate to (and potentially help sustain) CNS inflammation in neuroinflammatory conditions. Conceptually, immune signals and immune cells can influence brain physiology through BBB trafficking, meningeal/CSF interfaces, and cytokine-mediated communication, providing a framework in which peripheral transcriptional states may reflect neuroimmune activation even when CNS tissue is not directly sampled^54^.

In HTLV-1 disease specifically, prior work supports the presence of a compartmentalized intrathecal immune response, including the enrichment and accumulation of HTLV-1–specific cytotoxic T cells in CSF, which is consistent with ongoing immune surveillance and inflammatory effector activity within the CNS^55^. Moreover, CSF inflammatory mediators such as CXCL10 have been repeatedly associated with HAM/TSP neuroinflammatory activity and prognosis, suggesting that chemokine axes implicated in leukocyte recruitment may be operative in this setting^56^.

More broadly, IFN-γ–linked immune programs are increasingly recognized as capable of shaping microglial states and neural network function, offering a plausible mechanistic bridge between peripheral immune activation signatures and downstream neuroinflammatory consequences^57^. Taken together, these observations support a cautious interpretation that PBMCs transcriptomic remodeling in HTLV-1 may reflect systemic immune polarization coupled to CNS-relevant inflammatory pathways; definitive mechanistic linkage will require future studies integrating paired blood–CSF profiling, longitudinal sampling, and cellular trafficking/functional assays^58^

Our findings indicate that neuroimmune molecular networks are not unique to HTLV-1; similar signatures are detectable in leukocytes and are consistently altered across multiple conditions. In oncogenic viral hepatitis (HBV/HCV/HDV)^59^ and HPV lesions^60^, transcriptomic shifts in PBMCs highlight vesicle trafficking/endocytosis and neurotransmitter receptor pathways, with neuroimmune networks emerging as key discriminators. This is consistent with hybrid cytokine and synapse-like communication. Importantly, comparable coordinated neuroimmune dysregulation is also observed in non-viral conditions such as major depressive disorder, where shared blood–brain signatures (including PAX6 and related genes) support a common neuroimmune axis measurable in peripheral leukocytes^30^.

The emergence of neuron-like signaling programs in circulating leukocytes is biologically significant because it suggests that peripheral immune cells are not merely responding to inflammation but are transcriptionally primed to sense and integrate neuroactive cues (e.g., neurotransmitters/neuropeptides) and participate in synapse-like modes of intercellular communication, which are core principles of neuroimmune crosstalk^60,61^.

In this framework, the acquisition (or amplification) of neuronal/synaptic-associated modules in PBMCs may lower the barrier for bidirectional neuroimmune signaling by (i) increasing immune-cell responsiveness to neural mediators via neurotransmitter receptors and related signaling components^62^, (ii) promoting vesicle trafficking/exocytosis and contact-dependent communication that can shape cytokine release, antigen presentation, and migratory behavior^63^, and (iii) potentially enhancing the capacity of peripheral immune cells to influence neuronal circuits indirectly through cytokine-driven synaptic remodeling and plasticity^64^.

Importantly, analogous peripheral “reprogramming” is consistent with observations across other neuroinflammatory and neurodegenerative contexts. Blood transcriptomic studies in multiple sclerosis demonstrate that PBMCs gene-expression profiles capture disease-relevant programs and can stratify clinical stages^65^, while single-cell profiling of CSF immune cells reveals convergent neuroinflammatory transcriptional landscapes across diverse neurological disorders, supporting the idea of shared neuroimmune modules that transcend a single etiology^58^.

Likewise, growing evidence in Alzheimer’s disease highlights that peripheral immune cells and their trafficking into the CNS can amplify neuroinflammation and contribute to neuronal dysfunction, reinforcing the mechanistic plausibility that peripheral neuroimmune signatures are functionally coupled to CNS outcomes^66^. Together, these data support the interpretation that neuron-like signaling programs in leukocytes may represent an immunological “sensory” adaptation to neural microenvironments that becomes maladaptive when dysregulated, potentially altering neuroimmune communication and contributing to synaptic dysfunction through inflammatory mediators and neuromodulator pathways.

This study has limitations that should be considered when interpreting the results. Although we identify robust and reproducible transcriptional patterns associated with neuroimmune-related pathways in peripheral immune cells from HTLV-1–infected individuals, our conclusions are primarily based on transcriptomic analyses, pathway enrichment, and statistical associations.

Accordingly, the data presented here do not support the inference that peripheral immune cells acquire canonical neuronal functions, such as synaptic transmission or neurotransmitter release. Instead, our findings reflect the activation of molecular programs shared between immune and nervous system biology, many of which involve genes with well-established roles in T cells and other immune populations (e.g., *ATF4, ATP6V0D1, CXCR4*). The classification of these genes within neurobiological contexts should therefore be interpreted as evidence of functional overlap and pathway convergence, rather than proof of neuron-like identity or behavior.

We acknowledge that additional molecular, cellular, and mechanistic studies will be required to determine whether and how these transcriptional signatures translate into functional neuroimmune interactions and contribute causally to disease pathogenesis. However, such experiments are beyond the scope of the present work. Taken together, this study should be viewed as exploratory and hypothesis-generating, providing a systems-level framework for neuroimmune transcriptional remodeling in HTLV-1 infection. Our results establish a foundation for future experimental validation and longitudinal studies aimed at dissecting the mechanistic relevance of these neuroimmune signatures in disease progression.

In conclusion, this study provides systems-level evidence that HTLV-1 infection is accompanied by neuroimmune-related transcriptional remodeling, with the most robust and consistent alterations observed in ATL. Although HAM/TSP also exhibits changes in selected neuroimmune markers, our data indicate that the magnitude and direction of these signatures differ substantially between ATL and HAM/TSP, arguing for phenotype-specific neuroimmune programs rather than a single shared axis across HTLV-1–associated diseases. By integrating transcriptomic and flow cytometric analyses of PBMCs, we uncovered a consistent pattern of disrupted neurobiological gene programs, such as ATF4, CXCR4, and ATP6V0D1, which are already detectable in ACs and become progressively dysregulated with disease severity. These signatures suggest that alterations in neural-immune crosstalk are early events in HTLV-1 pathogenesis and may drive both malignant transformation and neuroinflammation.

Rather than treating ATL as purely oncological and HAM/TSP as exclusively neurological, our findings support a unified model of HTLV-1 pathology as a spectrum of distributed neuroimmune disintegration. PBMCs emerge as accessible sentinels of this systemic dysfunction, offering valuable insights into both peripheral and CNS-related mechanisms. This reconceptualization has important implications for the development of targeted diagnostics and therapies at the neuroimmune interface, moving beyond conventional paradigms toward a more holistic understanding of HTLV-1 disease biology.

## Acknowledgement Funding Support

We gratefully acknowledge financial support from the São Paulo Research Foundation (FAPESP) through the following grants: 2018/18886-9 to O.C.M.; 2020/16246-2, 40013/2025-3, 2023/07806-2 to I.S.F.; and 2024/08016-8 to A.L.N.; 2023/12268-0 to A.S.A.; 2023/14417-2 to J.N.U; 2025/07090-2 to FYNV; 2020/12017-9 to M.C.M.M; and 2018/14933-2 to H.N. We also thank the National Council for Scientific and Technological Development (CNPq), Brazil, for grant 140481/2025-7 to R.G.N; 140013/2025-3 to A.S.A, and 309482/2022-4 to O.C.M., and the Coordination for the Improvement of Higher Education Personnel (CAPES) for the following support: CAPES/PROEX grant 88887.967996/2024-00 to Y.L.G.C..; and 88887.699840/2022-00 to F.Y.N.V. This work was supported by the Japan Society for the Promotion of Science KAKENHI (grant number JP21H05051, to K. Kataoka) and Japan Agency for Medical Research and Development (25ck0106789h0003 25ck0106860h0003, and 25gm1810002h0004, to K. Kataoka).

## Conflict of Interest

The authors declare no conflict of interest.

## Authors’ Contributions

F.Y.N.V., C.M.S., M.D.V. and O.C.M. co-wrote the manuscript; F.Y.N.V, A.L.N., J.N.U., D.L.M.F., L.F.S., D.G.A.F., A.S.A., R.G.N., Y.L.G.C., H.F.N.V., H.D.D., R.D., I.S.F., M.D.V., and O.C.M. provided scientific insights; F.Y.N.V. performed the bioinformatic analyses; C.M.S. and M.D.V. conducted the flow cytometry experiments using human samples and performed the corresponding statistical analyses; Y.S., Y.K., J.K., and K.K. performed the single-cell experiments with human samples and provided the resulting datasets; M.C.C.M., A.S., and H.I.N. developed the Archipelago R package; M.D.V. supervised the protein expression analysis using flow cytometry, O.C.M. supervised the project.

## Notes

https://github.com/feryurinv23/HTLV-1-Induced-Neuroimmunome-Correlates-with-Disease-Progression-and-Severity

### Competing Interest Statement

The authors have declared no competing interest.

