## Supplementary data for "HTLV-1-Induced Neuroimmunome Correlates with Disease Progression and Severity"

[1] Laboratory of Psychoneuroimmunology, Selye Lab, University of São Paulo School of Medicine, São Paulo, Brazil.
[2] Department of Clinical and Toxicological Analyses, School of Pharmaceutical Sciences, University of São Paulo, São Paulo, Brazil.
[3] Imperial College London, Department of Infectious Diseases, Imperial College, UK.
[4] Interunit Postgraduate Program on Bioinformatics, University of São Paulo, São Paulo, Brazil.
[5] Department of Immunology, Institute of Biomedical Sciences, University of São Paulo, Brazil.
[6] Hospital Israelita Albert Einstein, São Paulo, Brazil.
[7] Institut Pasteur de São Paulo, São Paulo, Brazil.
[8] D'Or Institute for Research and Education, Rio de Janeiro, Brazil.
[9] Bioinformatics Multidisciplinary Environment, Federal University of Rio Grande do Norte, Brazil.
[10] Department of Medicine, Division of Molecular Medicine, Laboratory of Medical Investigation 29, University of São Paulo, Brazil.
[11] Ann Romney Center for Neurologic Diseases, Brigham and Women's Hospital, Harvard Medical School, Boston, MA, USA
[12] Division of Molecular Oncology, National Cancer Center Research Institute, Tokyo, Japan.
[13] Division of Gastroenterology and Hepatology, Department of Internal Medicine, Keio University School of Medicine, Tokyo, Japan.
[14] Division of Hematology, Department of Medicine, Keio University School of Medicine, Tokyo, Japan
*Contributed Equally

### Corresponding authors

Fernando Yuri Nery do Vale;

Otavio Cabral-Marques, MSc, PhD;

### Supplementary methods

We employed a multi-dimensional systems biology framework to investigate the molecular underpinnings of HTLV-1 pathogenesis (**Supplementary Figure 1**). Building on our single-cell RNA sequencing (scRNA-seq) dataset, which comprises 233,093 PBMCs, we aimed to validate and further explore the dysregulation of the neuroimmunome identified in this cohort. To achieve this, we integrated publicly available bulk RNA sequencing from PBMCs and microarray datasets (n = 200) derived from independent cohorts of individuals with HTLV-1 infection, including those with ATL, HAM/TSP, and asymptomatic carriers, as well as healthy controls. Our inclusion criteria focused on individuals infected with HTLV-1 and diagnosed with the associated diseases, specifically ATL, HAM/TSP, and asymptomatic carriers. Importantly, we excluded patients undergoing any form of treatment, individuals with co-infections, or those with other associated diseases to reduce confounding factors and ensure the specificity of the findings to HTLV-1 infection.

This integrative strategy enabled cross-platform validation of neuroimmune gene signatures and reinforced the consistency and generalizability of the observed molecular alterations across distinct biological contexts and transcriptomic resolutions. To further strengthen these findings, we also performed flow cytometric analysis to validate key neuroimmune markers at the protein level and assessed their association with HTLV-1 proviral load across clinical subgroups.

All statistical analyses were performed using R Studio software (The R Foundation for Statistical Computing) (v. 4.3.2). R packages from the

Bioconductor and CRAN repositories were utilized to analyze the transcriptomic data and create data visualizations[12]

**Data availability**

The tables used to generate and visualize the results of our study are provided as supplementary files. These tables include comprehensive data inputs for the analysis and detailed output results that encapsulate the findings and insights derived from our research. The supplementary input tables outline the dataset structure, variables, and relevant parameters, while the output tables display processed results, statistical metrics, and visual summaries that support our conclusions. The datasets used in the analysis were publicly available at the GEO datasets under the accession codes GSE29312, GSE38537, GSE233437, and GSE33615, as well as in the European Genome-phenome Archive (EGA) under the accession number EGAS00001004936.

The total number of individuals per dataset is as follows: 100 individuals from EGAS00001004936, 45 individuals from GSE29312, 20 individuals from GSE38537, 16 individuals from GSE233437, and 50 individuals from GSE33615. The distribution of individuals in each subtype is as follows: 41 individuals with Acute, 46 with Chronic, 6 with Smoldering, 27 Asymptomatic, 38 Healthy, 6 with Lymphoma, and 18 with HAM/TSP. The custom code used for all analyses is available at https://github.com/feryurinv23.

**Single cell analysis**

scRNA-seq data from 30 ATL patients, 11 HTLV-1 asymptomatic carriers (ACs), and 4 healthy donors were analyzed. The Seurat object was loaded into RStudio using the Seurat package[1–3] for further processing. Quality control measures were applied to ensure high data integrity. Cells with fewer than 200 or more than 2,500 detected RNA features were excluded to eliminate low-quality cells, such as dead or dying cells with minimal transcriptional activity, and potential doublets with abnormally high RNA content. Additionally, cells in which more than 5% of unique molecular identifiers (UMIs) were derived from mitochondrial genes were removed to avoid artifacts associated with cellular stress or apoptosis. The data were then normalized and scaled to correct for variations between cells, enabling more accurate comparisons.

Dimensionality reduction was performed using principal component analysis (PCA), followed by cell clustering via the Louvain algorithm. The resulting structure was visualized using Uniform Manifold Approximation and Projection (UMAP), as previously described by Koya et al. (2021)[4]. Cell types, including malignant and non-malignant cells, were assigned based on previous reports by Koya et al., 2021[4], which utilized T-cell marker expression, TCR repertoire analysis, and mRNA/ADT levels of canonical lineage markers[4]. To confirm these classifications, we applied the FeaturePlot() function using gene markers listed in **Supplementary Table 01**.

For downstream analyses, differentially expressed genes (DEGs) were identified using the FindMarkers() function, comparing gene expression profiles between ACs, ATL subtypes, and the healthy control group.

**Functional enrichment, clustering, and interactome**

Gene Ontology (GO) enrichment analysis was performed using the clusterProfiler package[5–7] in R (v. 4.3.2) and the EnrichR[8–10] bioinformatics tool to identify enriched biological processes (BPs). Pathway analysis was conducted using the KEGG database[11], with a significance threshold of adjusted *p*-value ≤ 0.05 applied across all datasets. To evaluate the nervous and immune clusters formed by BPs, we utilized the Appyters[12] web tool, integrated with EnrichR[12], was utilized. This approach facilitated the identification of enriched BPs, which were then grouped into clusters and visualized using a scatter plot. The Leiden algorithm was applied for clustering[13], with dimensionality reduction performed using Uniform Manifold Approximation and Projection (UMAP). The results were visualized using the ggplot2 package[14] in R (v. 4.3.2). Additionally, a network analysis of genes associated with enriched BPs was conducted to explore the neuroimmunological interactome, achieved using the cnetplot() function from the enrichplot package[15] in R (v. 4.3.2).

**Characterization of neuroimmunome genes and terms related to biological processes**

To characterize DEGs involved in the neuroimmune response, we utilized the [16]BPs database to identify genes annotated as belonging to both the immune and nervous systems. In the subsequent step, we selected BP terms for dot plot and network visualization. For neurological processes, we used the following keywords: “synapse”, “synaptic”, “neuro”, “glia”, “nervous”, “axon”, “glutamate”, “gliogenesis”, and “nerve.” For immune-related processes, we included: “response to virus”, “immune response”, “B cell”, “T cell”, “cytokine”, “leukocytes”, “humoral immune”, “lymphocytes”, “immunoglobulin”, “interferon”, “innate immune", and "adaptive immune." This enrichment analysis provided deeper insight into how these neuroimmune genes may contribute to the biological functions of both systems and their relevance in the context of our study.

**Assessing Synaptic-related Processes at Single-Cell Level**

We applied a comprehensive workflow to identify pathways and BPs related to synaptic activity and the neuro-immune process, aiming to gain new insights into the role of peripheral neuroimmune communication at the single-cell level in the context of HTLV-1. By integrating different bioinformatics approaches, we explored the peripheral synaptic gene expression associated with DEGs in PBMCs from individuals with HTLV-1 infection, contributing to our hypothesis on neuroimmune interactions.

[14,1718191120]

*Neurotransmission-linked gene networks associated to disease stage*

Additionally, neurotransmitter pathways were analyzed to investigate the DEGs identified in individuals with HTLV-1 infection in PBMCs associated with the ACs and ATL subtypes. The aim was to understand the role of neurotransmitter-related gene expression and the neurotransmitter families to which they belong. This analysis was conducted using the KEGG database[11], combined with network visualization via ggplot2[14,16] to illustrate functional interactions[18].

*Cell-enriched Island using ArchipelaGO*

We applied the workflow to identify cell-enriched "islands" associated with specific BPs, using UMAP components as spatial coordinates. This package integrates a gene expression matrix, spatial cell coordinates from UMAP, and user-defined gene sets to identify spatially enriched regions, termed "islands". These islands represent clusters of cells with high enrichment for specific biological processes. We focused on the top five BP gene sets related to neuroimmune interactions, obtained using the annotationDBi[17] and org.Hs.eg.db[18] packages from Bioconductor.

Initially, enrichment scores for each cell were calculated using over-representation analysis (ORA), with Fisher's exact test employed to determine statistical significance. The p-values were transformed into enrichment scores defined as -log(p). These scores were then used to generate spatial enrichment landscapes via kernel density estimation, where UMAP components and their respective scores were integrated to create a continuous topographical map. Islands were identified by applying a user-defined threshold ("water level") to the topographical map, delineating regions of significant enrichment. Each island was further characterized by its cellular composition, determined based on the cell types present within its polygonal contours.

*Enrichment analysis of synaptic and disease processes*

Human KEGG pathway annotations were retrieved in R (v. 4.3.2) using the KEGGREST package and curated within the tidyverse framework. To focus on neurotransmission-related signatures, pathways were selected through keyword filtering of pathway names, retaining entries containing ‘dopamin’, ‘gaba’, ‘glutamat’, ‘seroton’, or ‘cholin’. Enrichment analysis for this step was conducted using only upregulated genes. Additionally, disease-associated enrichment was assessed using the Jensen DISEASES Curated 2025[19] in EnrichR plataform[8], considering the full set of DEGs for each comparison.

**Metanalysis of transcriptomic data.**

Eligible studies were identified using the Gene Expression Omnibus (GEO)[20] by searching for peer-reviewed datasets indexed with the terms "HTLV-1," "Homo sapiens," and expression profiles from “high throughput sequencing” (bulk or single-cell RNA-seq) and “expression profiling by array.” Studies had to contain healthy controls and include a minimum of 8 total samples. Initially, 64 studies reporting transcriptional data from human subjects were identified (**Fig. 4**).

Studies were excluded based on the following criteria: (1) therapeutic-focused analyses (n = 12); (2) use of cultured cell lines, tissues, or genetically modified viruses (n = 30); (3) fewer than three infected samples (n = 11); (4) duplicated or replicated samples (n = 3); (5) time-course or longitudinal datasets (n = 3); and (6) super series datasets (n = 1). After applying the exclusion criteria, we incorporated two datasets provided by our collaborators, resulting in a total of five datasets deemed eligible for inclusion in our meta-analysis workflow, along with one additional single-cell dataset.

These datasets included a total of 200 participants: 135 with ATL, 47 ACs, 18 with HAM/TSP, and 45 healthy controls (**Fig. 4**). To comprehensively evaluate the transcriptional overlap between DEGs from scRNA-seq of 233,093 PBMCs and the metaDEGs (obtained through meta-analysis as described below), further analyses were conducted.

The meta-analysis was conducted using the web tool "ExpressAnalyst"[21,22] to quantitatively synthesize results from multiple studies on gene expression. Specifically, the datasets GSE29312, GSE38537, GSE233437, GSE33615, and EGAD000010007024 (bulk RNAseq) were utilized for the meta-analysis under the conditions asymptomatic, ATL, and HAM/TSP. Metadata for each dataset, including clinical staging (subtype), sex, age, and other available clinical variables, were carefully reviewed and harmonized to maximize cross-cohort consistency and ensure greater dataset homogeneity for downstream comparative analyses. All metadata can be accessed via the corresponding accession code for each dataset.

Data annotation involved specifying the value type (raw data), data type (microarray or bulk RNAseq), selecting the human organism, and specifying the ID type for all datasets. Missing values were addressed by excluding features with more than 50% missing values and estimating the remaining missing values using KNN imputation (feature-wise). Subsequently, filtering and normalization were performed using a 15% variance filter based on the inter quantile range and a 5% abundance filter (relative percentile). Batch effects across transcriptomic datasets were corrected using ComBat prior to integrative analyses. Expression values were adjusted by modeling “dataset/study” as the batch factor while including the relevant biological condition(s) of interest as covariates in the design matrix to ensure the preservation of true biological signals. ComBat applies an empirical Bayes approach to estimate gene-wise location and scale shifts associated with each batch removing these systematic, non-biological differences and harmonizing expression values across datasets. The effectiveness of batch correction was evaluated by inspecting expression density distributions and principal component analysis (PCA) before and after adjustment.

Differential expression analysis was conducted using the Limma framework. For each dataset, gene-wise linear models were fitted to normalized, log-transformed expression values, and the relevant group comparisons were specified through contrasts. Limma's empirical Bayes moderation was applied to shrink gene-specific variance estimates toward a pooled value, enhencing robustness and power, particularly in studies with modest sample sizes.

P-values were adjusted for multiple testing using the Benjamini–Hochberg false discovery rate (FDR). Evidence for differential expression across datasets was integrated by combining adjusted p-values with Fisher's method. Genes were considered significantly differentially expressed if they met an adjusted p-value (FDR) < 0.05 and exhibited a non-zero average fold change (average fold change ≠ 0.0).

**Principal component analysis.**

PCA was performed using spectral decomposition with the factoextra[23] and FactoMineR packages[24] in R (v. 4.3.2). Eigenvalues were calculated based on the contributions of neuroimmune genes associated with BPs to assess their influence on PCA. Eigenvalues greater than one were considered critical for determining group segregation[25]. The first two principal components were analyzed to evaluate group separation and the role of neuroimmune genes in distinguishing healthy controls from ATL conditions. The top 20 genes contributing most to the first two dimensions were selected for further analysis.

**Relative effect analysis**

We evaluated the relative effects of the highest neuroimmune genes identified by PCA (top 20 genes) in ATL compared to a healthy control group using a Multivariate Analysis of Variance (MANOVA) with a bootstrap[26] approach for enhanced reliability, involving 1,000 re-samplings. This method allowed us to account for variability in the data and calculate confidence intervals (CIs) for the relative effects, providing a robust measure of association. The statistical analysis was performed using the R packages npmv[27] and reshape[28].

**Gradient boosting machine**

We employed the Gradient Boosting Machine (GBM) method based on decision tree methodology using the gbm R package[29] for feature selection to identify the most relevant genes contributing to the disease. This approach helped reduce the dimensionality of genomic data and focus on the most informative genes[30]. The model was performed using the top 20 high-score neuroimmune genes identified by PCA on 67 samples from GSE36516 (46 samples in the ATL group and 21 samples in the control group). A total of 500 trees were used in the analysis, and the Huberized distribution method was applied. The partial effects of the genes analyzed in the GBM model were examined using all the trees in the model and a grid resolution of 100 points.

**Additional data visualization features**

To better understand the differences in gene expression between the healthy control group and the ATL condition, we visualized gene expression using violin plots generated with the ggplot2[16] package. Statistical differences between groups were assessed using the Wilcoxon test, implemented through the stat_compare_means function from the ggpubr package[31].

**Ethical Study Approval**

Ethical approval for the collection of blood samples from healthy individuals has been granted by the Imperial College Research Ethics Committee (ICREC reference number 6931613). Ethical approval for blood collection from individuals with HTLV-1 infection has been granted by the Health Research Authority UK (South Central – Oxford C Research Ethics Committee, REC reference: 20/SC/0226). The dataset **EGAS00001004936** was generated by **Koya et al. (2021)**[4] as part of studies approved by the Institutional Review Board of the National Cancer Center and participating institutions. The data are deposited in the European Genome-phenome Archive (EGA**)** and are available under controlled access to facilitate secure data sharing. Patient samples were collected at the University of Miyazaki Hospital and Imamura General Hospital. All participants provided written informed consent in accordance with the Declaration of Helsinki, and the study protocols were approved by the relevant institutional review boards.

**Single cell experimental analysis**

Single-cell experiments were performed and analyzed by our collaborators in Japan, who originally published these single-cell datasets. Detailed descriptions of cohort composition, ethical approvals, mouse generation, and experimental procedures are provided in **Koya et al., 2021**[4].

*Human Samples*

Primary PBMCs were obtained from 11 ACs (11 samples) and 30 ATL patients (34 samples, including four sequential samples: 19 acute, 12 chronic, and 3 smoldering subtypes), along with samples from 3 healthy donors. All patients were untreated with systemic chemotherapy at the time of peripheral blood (PB) collection, except for one patient with sequential sampling who received chemotherapy between the first (ATL01) and second (ATL01-2) PB collections. HD samples were purchased from Zen-Bio, Precision Bioservices, iQ Biosciences, and HemaCare Corporation. The available clinical characteristics and sequencing information are summarized in Koya et al. (2021)[4].

Asymptomatic carriers (AC) are HTLV-1–positive individuals with no clinical disease and no evidence of malignant T-cell expansion[32]. Clinical classification of ATL, which is divided into indolent types (chronic, smoldering) and aggressive types (acute, lymphoma), rather than a strict chronological sequence, acute ATL is not necessarily a stage that comes after chronic ATL, but a distinct aggressive subtype[33].

*Sample Preparation, Library Preparation, and Sequencing*

PBMCs were isolated from whole blood using Ficoll-Paque PLUS (Cytiva) and cryopreserved in CELLBANKER 1plus (TaKaRa Bio) at −150°C, following the manufacturers' instructions. Frozen samples were rapidly thawed in prewarmed RPMI-1640 supplemented with 10% FBS, washed by centrifugation, resuspended, filtered through a 70-μm nylon mesh, and stained with DAPI for live/dead discrimination. Cells were sorted by gating on forward/side scatter, excluding doublets, and selecting DAPI^low viable cells. After sorting 1 × 10^6 live cells, pellets were centrifuged (300 × g, 4°C, 7 min), resuspended in Cell Staining Buffer with Human TruStain FcX, and incubated on ice for 10 min.

For CITE-seq profiling, PBMCs were stained with TotalSeq-C antibody panels (BioLegend): a 99-antibody set for 18 samples and a 105-antibody set for 31 samples (each antibody at 0.2 µg). scRNA-seq, scADT-seq, and scTCR/BCR-seq libraries were generated using 10x Genomics Chromium Single-Cell 5′ and V(D)J workflows (v1/v1.1), including Feature Barcode libraries for cell-surface proteins, following the manufacturer’s protocol. Libraries were sequenced as 150-bp paired-end reads primarily on an Illumina NovaSeq 6000 (S4 kit) at Macrogen; three scBCR-seq libraries (ATL12, ATL14, ATL22) were sequenced in-house on an Illumina NextSeq 500 using the High Output kit.

**Flow cytometric analysis**

Cryopreserved PBMCs from healthy individuals, ACs, and people with ATL, or HAM/TSP were prepared and stained for analysis. First, the PBMCs were thawed in RPMI 1640 medium supplemented with human AB serum and Benzonase. After centrifugation and resuspension in RPMI 1640 medium supplemented with 2mM L-Glutamine, 5% human Serum AB and 1x Pen/Strep, the cells were rested at 37°C for 2 hours. The cells were first stained with Fixable Blue viability dye, followed by Fc receptor blocking and a 30-minute incubation with a chemokine receptor antibody for CXCR4 (12G5, Biolegend) at 37°C. They were further stained for surface markers: CD3 (UCHT1, Biolegend), CD4 (S3.5, Thermofisher), CD45RA (HI100, Biolegend), CD45RO (UCHL1, Biolegend), CD16 (3G8 RUO, BD), CD14 (M5E2 RUO, BD) and CD20 (2H7, Biolegend); and fixed using the FoxP3 fixation and staining buffers (Thermo Scientific). After fixation, the cells underwent a permeabilization step with Perm/Wash solution, followed by intracellular blocking and intracellular staining using antibodies targeting SKIL (OTI3E2, Novus Biologicals), ATF4 (S360A-24, Novus Biologicals), V-ATPase D1 (*ATP6V0D1*; D-4, Santa Cruz Biotechnology), Senataxin (*SETX;* 4G1, Novus Biologicals), PTBP1 (3D6D8, Thermofisher), VAMP2 (EPR12790, Abcam), and Ku80 (*XRCC5;* 2G5E7, Thermofisher). This staining pipeline was previously established and validated by our collaborators[34–36]. TIAM2 was excluded due to the unavailability of a validated anti-TIAM2 antibody.

### Supplementary Figures

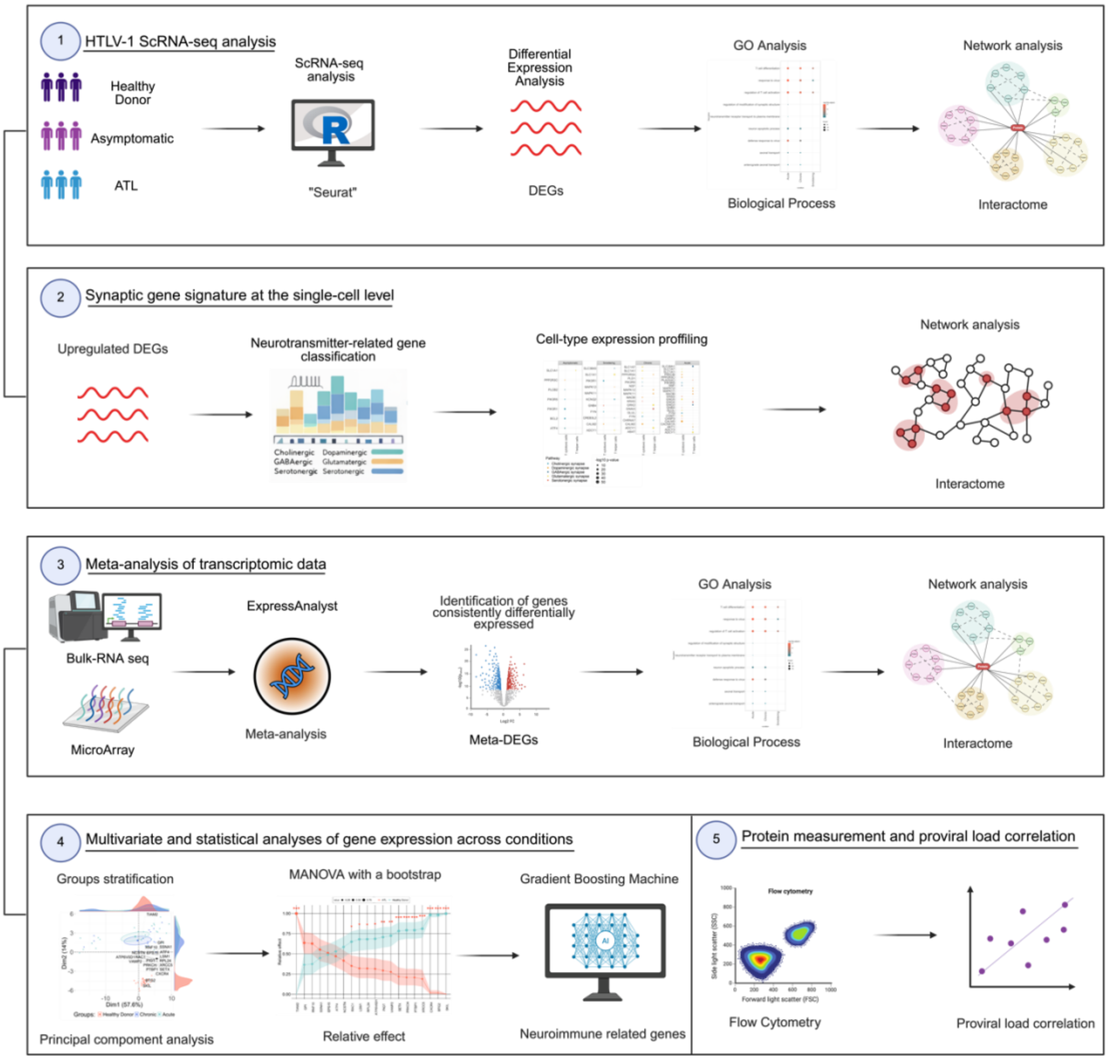


**Supplementary Figure 01: Systems biology workflow illustrates the integrative analytical pipeline used in this study as described in the Materials and Methods.**

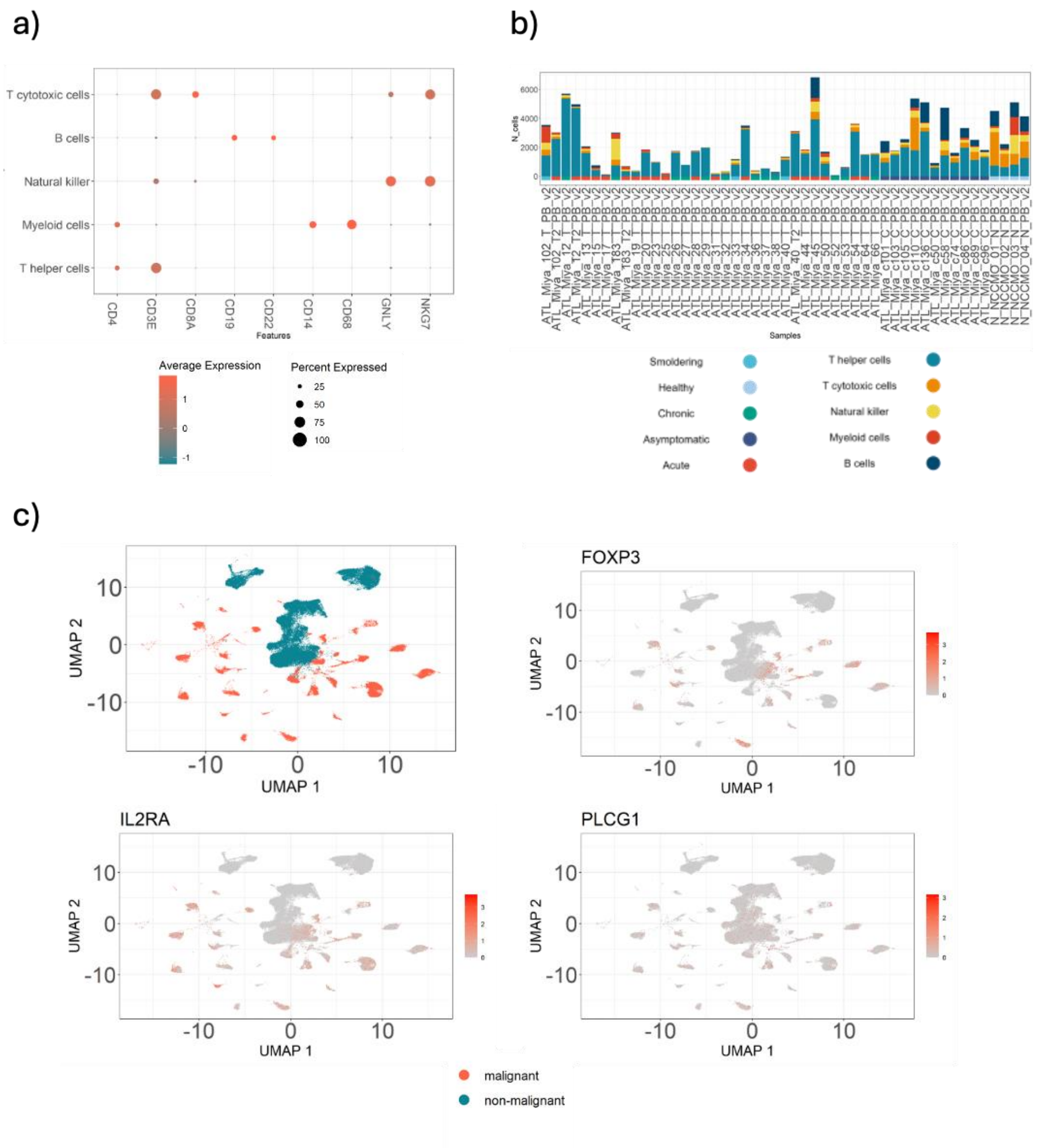


**Supplementary Figure 2: Neuro-Immune Gene Expression Profiles Associated with Malignant Cells in Single-Cell Data. (a)** Dot plot showing the expression of selected immune markers across major immune cell types, highlighting the specificity of genes such as *CD3E*, *CD4*, *CD14*, *CD19*, *CD68*, and *NKG7*. (b) Bar plot showing the total number of captured cells per individual sample. Bars are partitioned by clinical condition, enabling comparison of both absolute cell yields and relative condition contributions across the cohort. The plot highlights substantial inter-sample variability in cell recovery and differences in condition representation between samples. (c) UMAP feature plots displaying the expression of selected gene markers (*FOXP3*, *IL2RA*, *PLCG1*), indicating their association with malignant cell clusters.

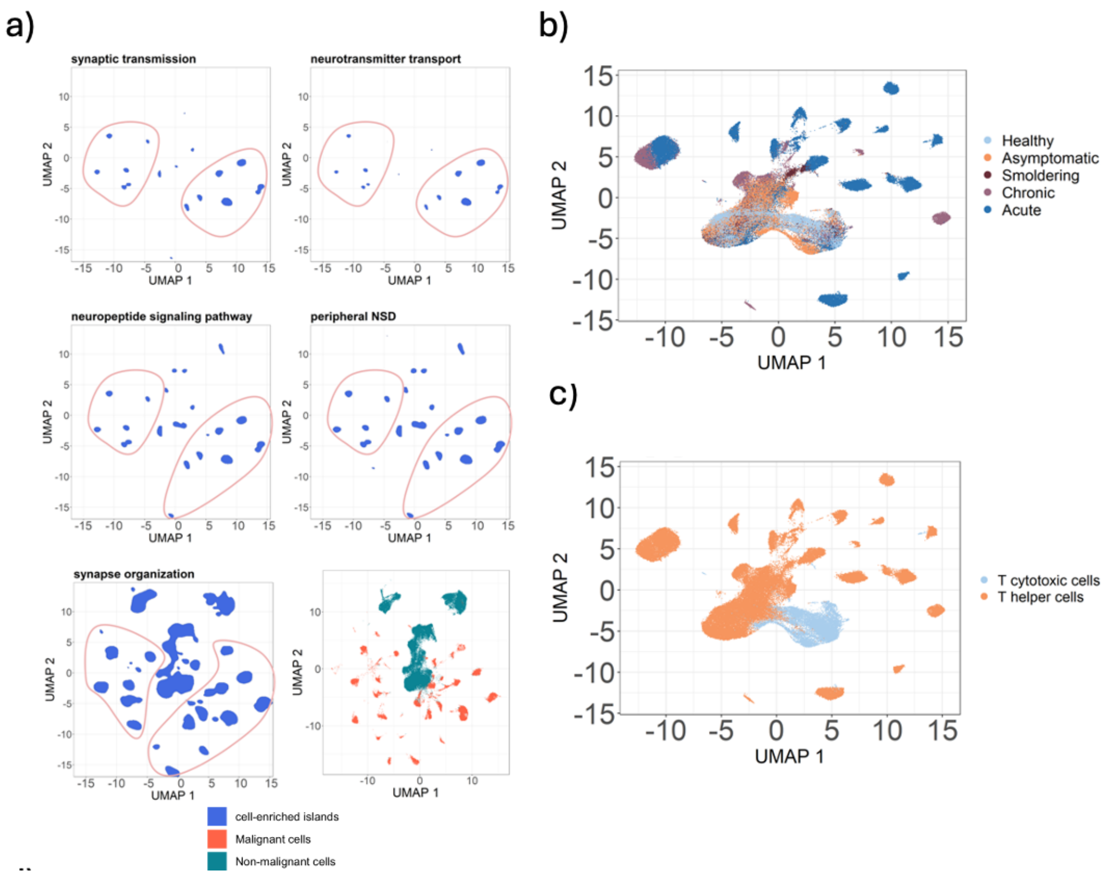


**Supplementary Figure 3: Neuroimmune-associated biological processes and synapse-related modules in single-cell T cells:** (a) Identification of cell-enriched "islands" associated with neuroimmune-related biological processes using UMAP embeddings as spatial coordinates. Gene expression matrices, UMAP-derived cell positions, and curated gene sets were integrated to detect spatially enriched regions representing clusters of cells with high BP activity. (b) UMAP projection of single-cell transcriptomes colored by clinical status (healthy, asymptomatic, smoldering, chronic, and acute). (c) UMAP projection highlighting T cell identities, distinguishing cytotoxic and helper T cell populations.

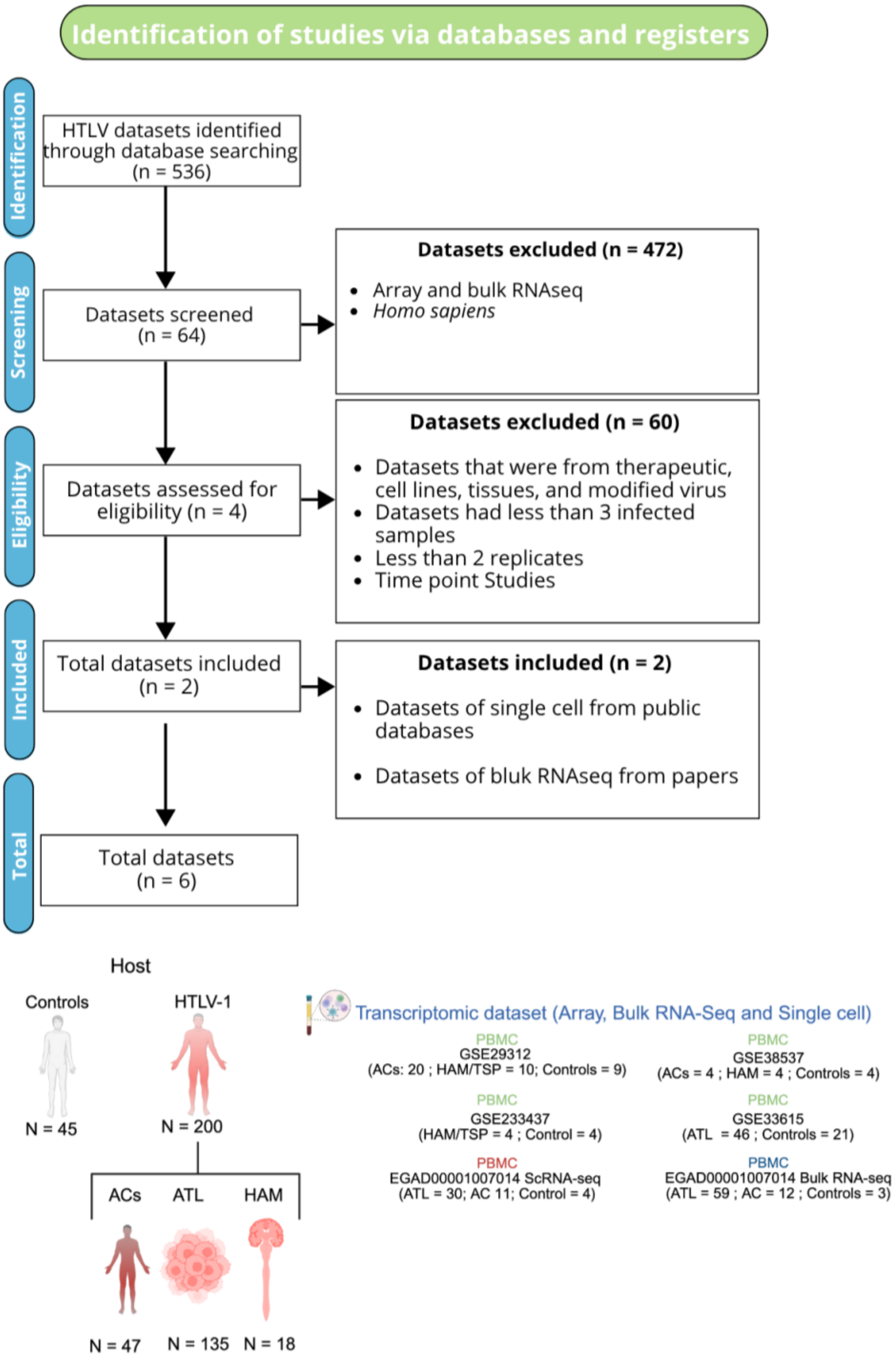


**Supplementary Figure 4: PRISMA workflow for dataset selection and meta-**

**analysis of RNA-seq and single-cell studies.** Flow diagram summarizing the selection process of transcriptomic datasets involving HTLV-1 infection. Out of 536 datasets identified through database searches, 64 datasets were screened after excluding non-human or array/bulk RNA-seq datasets. Following eligibility assessment, 60 datasets were excluded based on criteria such as the use of therapeutic samples, insufficient infected sample count, lack of replicates, or time-point designs. Ultimately, 6 datasets were included, comprising both single-cell and bulk RNA-seq data from public databases and published studies. The bottom section provides a breakdown of sample distribution across conditions (Controls, ACs, ATL, and HAM/TSP) and the associated transcriptomic datasets used.

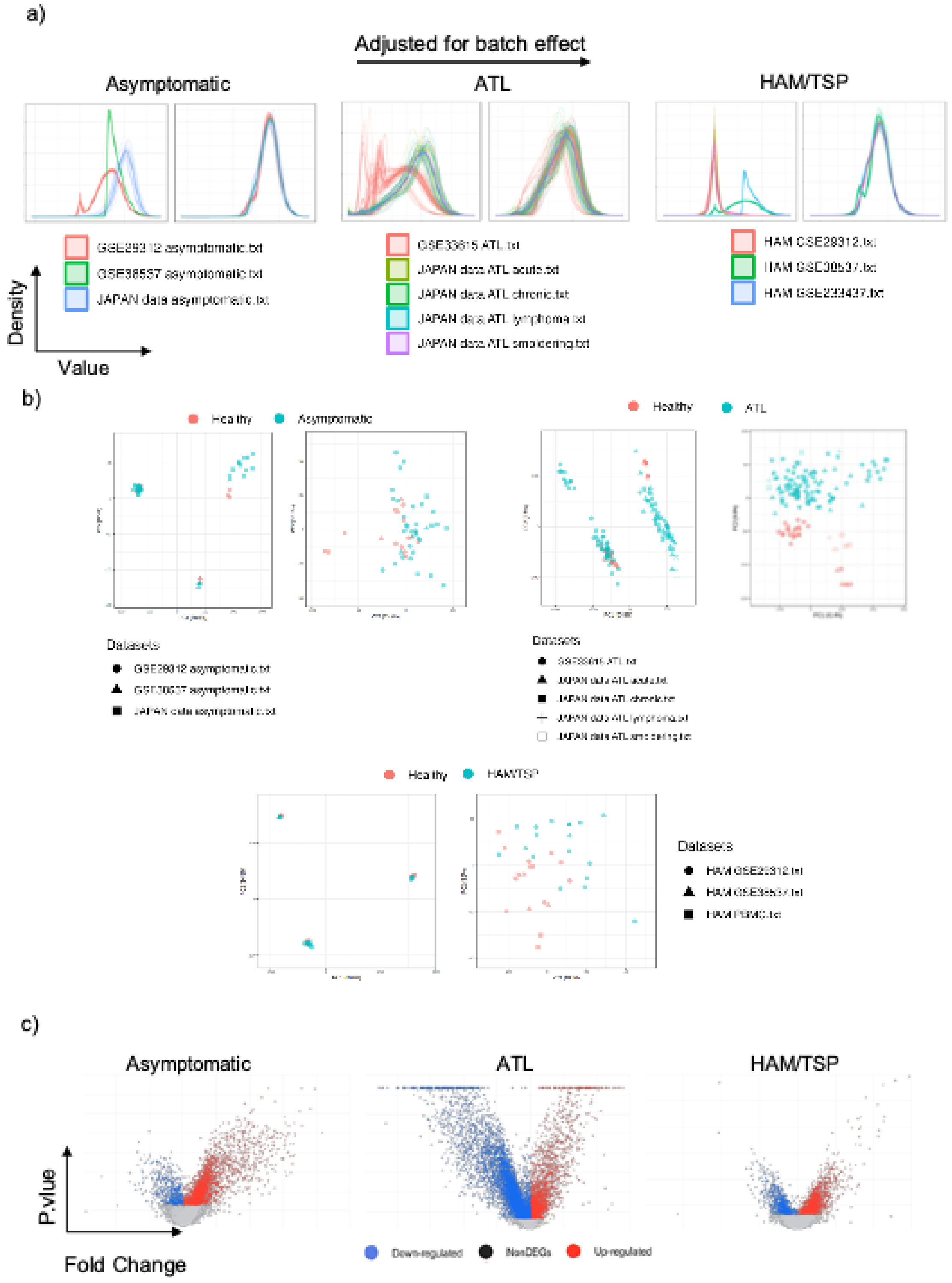


**Supplementary Figure 5: Meta-Analysis Reveals Transcriptional Patterns Across Multiple Conditions.** (a) Density plots showing distributions before and after batch effect correction, highlighting the harmonization of expression profiles across datasets and conditions. (b) Principal Component Analysis (PCA) plots illustrating the clustering of samples before (left) and after (right) batch effect correction, demonstrating improved data integration across different cohorts and disease stages. (c) Volcano plots of differentially expressed genes (DEGs) for each condition analyzed, showing significantly upregulated (red) and downregulated (blue) genes (adjusted $p < 0.05$).

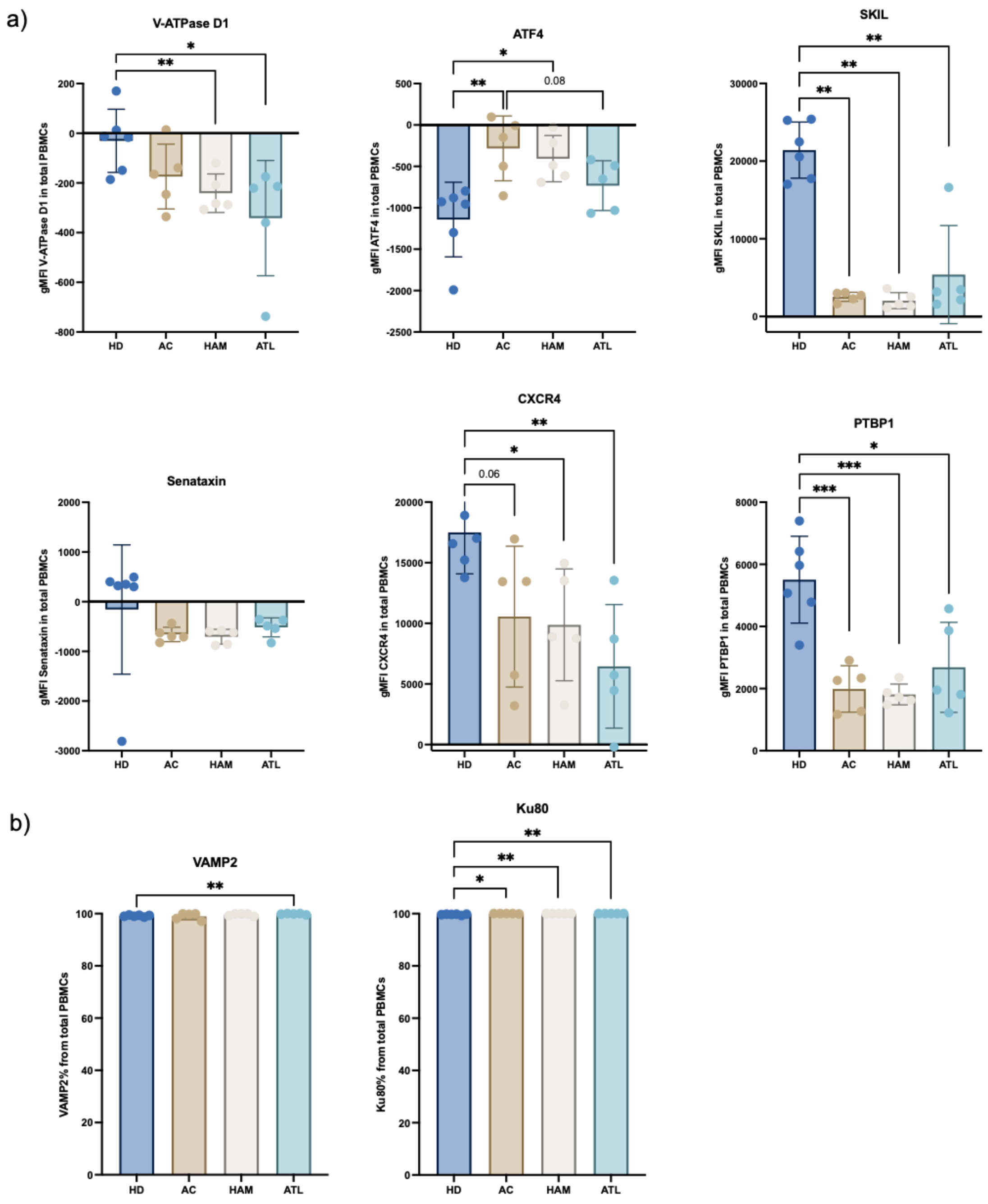

**Supplementary Figure 5: Intracellular protein expression in PBMCs from healthy individuals and individuals with HTLV-1 infection.** (a) Geometric mean fluorescence intensity (gMFI) of intracellular **V-ATPase D1, ATF4, SKIL, Senataxin (*SETX*), PTBP1** and **CXCR4**, measured by multiparameter flow cytometry in PBMCs from healthy donors (HD), ACs, and patients with HAM/TSP or ATL. (b) Frequency of total PBMCs expressing intracellular **VAMP2** and **Ku80** ***(XRCC5)***. Both proteins were ubiquitously expressed (>95% of PBMCs). Data points represent individual donors, with bars showing mean ± SD. Statistical significance was determined using unpaired t-tests with Welch's correction. $p < 0.05$ (*), $p < 0.01$ (**), $p < 0.001$ (***), $p < 0.0001$ (****).

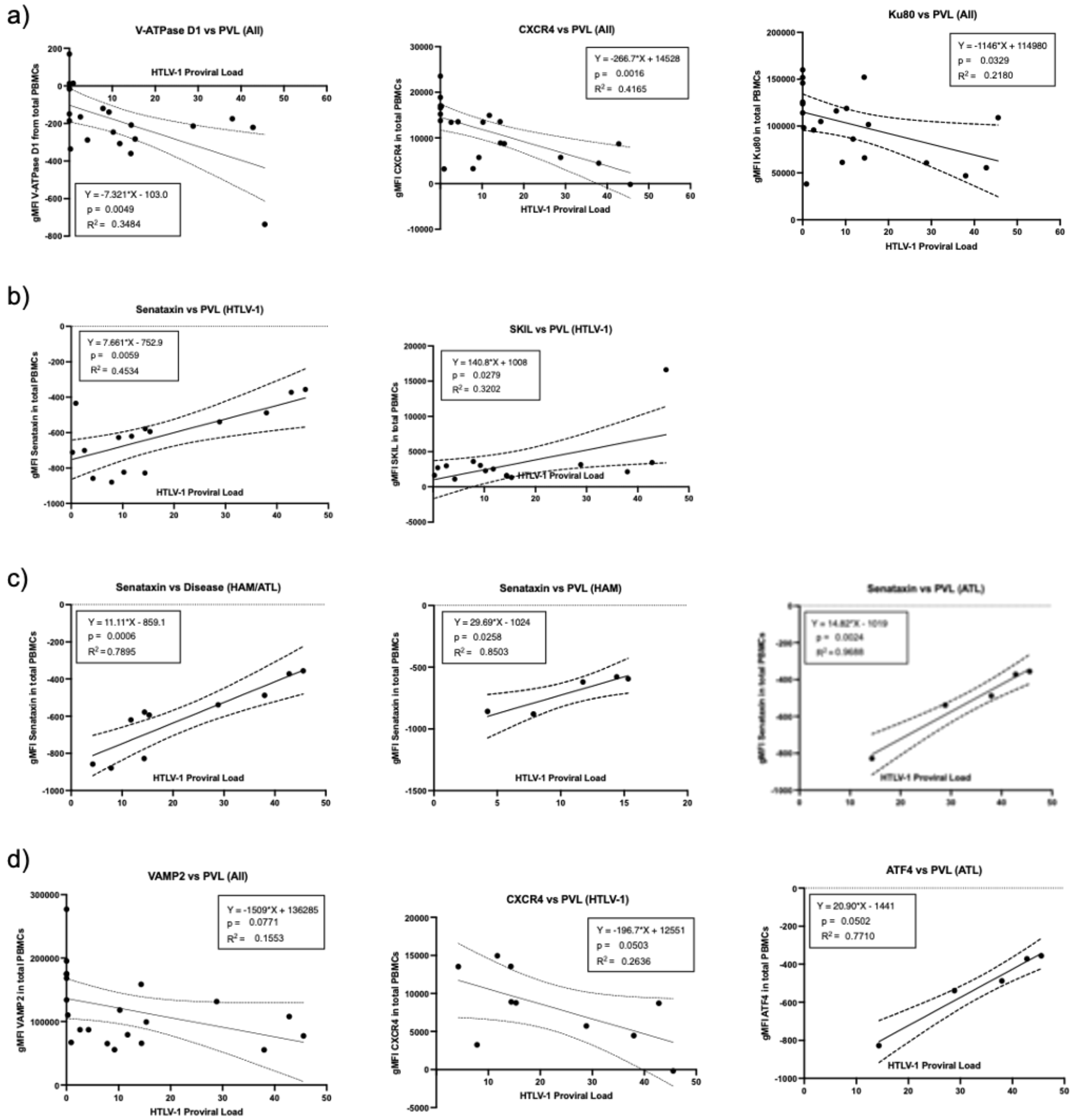


**Supplementary Figure 6: Correlation analysis between HTLV-1 proviral load and intracellular protein expression in PBMCs across disease subgroups.** Linear regression plots depicting the correlation between HTLV-1 proviral load (PVL) and the geometric mean fluorescence intensity (gMFI) of selected intracellular proteins in total PBMCs across **(a)** all individuals regardless of HTLV-1 infection, **(b)** individuals with HTLV-1 infection or **(c)** individuals with symptomatic HTLV-1 infection (HAM/ATL). **(d)** Correlations that did not reach statistical significance but show a trend towards it. Each point represents one subject; regression lines with 95% confidence intervals are shown.

platform for RNA-sequencing analysis in non-model species. Nat Commun *14*, 2995. https://doi.org/10.1038/s41467-023-38785-y.

27. Ewald, J., Zhou, G., Lu, Y., and Xia, J. (2023). Using ExpressAnalyst for Comprehensive Gene Expression Analysis in Model and Non-Model Organisms. Curr Protoc *3*. https://doi.org/10.1002/cpz1.922.
28. Kassambara, A., and Mundt, F. (2020). factoextra: Extract and Visualize the Results of Multivariate Data Analyses. Preprint.
29. Lê, S., Josse, J., and Husson, F. (2008). **FactoMineR** : An *R* Package for Multivariate Analysis. J Stat Softw *25*. https://doi.org/10.18637/jss.v025.i01.
30. Kaiser, H.F. (1960). The Application of Electronic Computers to Factor Analysis. Educ Psychol Meas *20*, 141–151. https://doi.org/10.1177/001316446002000116.
31. Xu, L.-W. (2015). Parametric bootstrap approaches for two-way MANOVA with unequal cell sizes and unequal cell covariance matrices. J Multivar Anal *133*, 291–303. https://doi.org/10.1016/j.jmva.2014.09.015.
32. Burchett, W.W., Ellis, A.R., Harrar, S.W., and Bathke, A.C. (2017). Nonparametric Inference for Multivariate Data: The *R* Package **npmv**. J Stat Softw *76*. https://doi.org/10.18637/jss.v076.i04.
33. Wickham, H. (2007). Reshaping Data with the **reshape** Package. J Stat Softw *21*. https://doi.org/10.18637/jss.v021.i12.
34. Ridgeway, G., and Developers, G. (2003). gbm: Generalized Boosted Regression Models. Preprint, https://doi.org/10.32614/CRAN.package.gbm https://doi.org/10.32614/CRAN.package.gbm.
35. Boldini, D., Grisoni, F., Kuhn, D., Friedrich, L., and Sieber, S.A. (2023). Practical guidelines for the use of gradient boosting for molecular property prediction. J Cheminform *15*, 73. https://doi.org/10.1186/s13321-023-00743-7.
36. Kassambara, A. (2016). ggpubr: “ggplot2” Based Publication Ready Plots. Preprint, https://doi.org/10.32614/CRAN.package.ggpubr https://doi.org/10.32614/CRAN.package.ggpubr.
37. Jones, E.M., Sourij, C., Stradner, M., Schlenke, P., Sereban, N., Moser, O., Quinlan, R., Short, C.-E., Harris, B.H.L., Fertleman, M., et al. (2025). Accelerated memory T cell decline and tolerogenic recall responses to SARS-CoV-2 vaccination in diabetes. Preprint, https://doi.org/10.1101/2025.02.08.637262 https://doi.org/10.1101/2025.02.08.637262.
38. Coulombeau, R., Selck, C., Giang, N., Al-Mohammad, A., Ng, N., Maher, A.K., Argüello, R., Scalfari, A., Varley, J., Nicholas, R., et al. (2025). <scp>Sphingosine</scp> -1-Phosphate Signalling Inhibition Suppresses Th1-Like Treg Generation by Reversing Mitochondrial Uncoupling. Immunology *174*, 153–166. https://doi.org/10.1111/imm.13870.
39. Maher, A.K., Aristodemou, A., Giang, N., Tanaka, Y., Bangham, C.R.M., Taylor, G.P., and Dominguez-Villar, M. (2024). HTLV-1 induces an inflammatory CD4+CD8+ T cell population in HTLV-1–associated myelopathy. JCI Insight *9*. https://doi.org/10.1172/jci.insight.173738.
